# Omics analysis reveals striking effects of progesterone receptor on mitochondria and mitochondria-mediated apoptosis independent of caspases in Breast Cancer cells

**DOI:** 10.1101/2025.03.06.641096

**Authors:** Qian Yee Woo, Pheck Khee Lau, Bernett Lee Teck Kwong, Natasa Bajalovic, Shi Hao Lee, Kye Siong Leong, Kai Yee Pow, Simra Hanan, Wei Meng, Soak Kuan Lai, Valerie CL Lin

## Abstract

The role of progesterone receptor (PR) in breast cancer remains controversial with conflicting reports from clinical and laboratory studies. To address these discrepancies, we conducted an integrated omics analysis of effects of agonist-activated PR in MCF-7 cells with elevated PR expression. PR agonist R5020 exerted strong antiproliferative and proapoptotic effects in these cells. Quantitative proteomics identified 4,915 PR-regulated proteins and 678 phosphorylated peptides, with nearly 100% verifiable rate by Western blotting analysis. The proteomics data was closely correlated with transcriptomic data. Key pathways upregulated included hypoxia, p53 signalling, TNFA signalling via NFKB, epithelial-mesenchymal transition, and KRAS signalling, while E2F targets, G2/M checkpoint, and mitotic spindle assembly were downregulated. R5020 broadly suppressed cell cycle regulators, including CDKs, cyclins, DNA replication proteins, and all components of the Ndc80 complex and chromosomal passenger complexes. Concurrently, it elicited significant changes in 200 mitochondrial proteins, upregulating many proapoptotic factors (e.g., BNIP3, NIX, AIF/AIFM1, AIFM2, ENDOG, HtrA2/Omi, Smac/DIABLO) and downregulating anti-apoptotic proteins (BCL-2, BCL-XL). This culminated in mitochondria-mediated apoptosis independent of effector caspases. The omics analysis also detected previously reported upregulation of pro-growth proteins such as EGFR, IRS2, and CCND1, but the upregulation was functionally futile and inhibitory phosphorylation of IRS2 at S306 increased 4-fold. In conclusion, this omics study achieved to date the most comprehensive and holistic understanding of PR-regulated proteins and molecular networks that are strongly anti-proliferative and proapoptotic with significant involvement of mitochondria. These findings suggest that pure PR agonists warrant evaluation as first-line endocrine therapy for breast cancer with high PR expression.

## Introduction

Estrogen receptor α (ERα) and progesterone receptor (PR) are expressed in two thirds of all primary breast cancer cases. ERα targeting therapies are the first line of endocrine therapy that are initially effective in the majority of the ERα and PR positive cases. However, 40% of patients are expected to experience recurrence within 20 years from diagnosis [1, 2]. ER-positive but PR-negative breast cancers have high recurrence risks [3]. Clinical trials in the 1980s and 1990s reported significant benefits of progestins (medroxyprogesterone acetate or megesterol acetate) as the second line endocrine therapy for women with recurrent and advanced breast cancer [4-7]. However, progestin therapy becomes controversial following reports of large trial results of hormone replacement therapy (HRT), which showed significantly higher incidences of breast cancer in women receiving estrogen and medroxyprogesterone acetate (MPA), compared to those receiving estrogen alone [8-11]. It has been suggested that the increased risk for breast cancer with MPA may be caused by crosstalk with androgen receptor signalling [12-14]. PR-specific synthetic progestins would be beneficial as a second line endocrine therapy.

In recent years, there has been a revival of interest in PR targeting therapies. Several clinical trials have been initiated to test PR targeting therapy with estrogen deprivation in early-stage primary breast cancers. PIONEER (NCT03306472) is a pre-operative window study of letrozole plus Megestrol Acetate versus letrozole alone in post-menopausal patients with ER-positive breast cancer. Progesterone is also being tested together with letrozole or tamoxifen in WinPro trial (NCT03906669) in early stage ERα+ and PR+ HER2-breast cancer. While the trial results are pending, these trials are largely inspired by the understanding of functional interplays between ERα and PR in progestin activated PR modulates ERα-regulated gene expression through reprograming ERα cistromes and redirects ER chromatin binding to achieve antiestrogenic signalling [15, 16]. This reprograming is associated with low tumorigenicity in progestin treated cells, and combination of selective PR modulator CDB4124 with anti-estrogen tamoxifen causes 70% tumor regression of T47D cell xenografts [16]. Thus, the advancements in the understanding of the molecular function of progestin and PR are crucially important in improving PR targeted therapy for breast cancer. Collectively, growing interests in targeting PR in breast cancer with various regimen reflect the importance of PR in breast cancer biology and the need to improve endocrine therapy.

Despite renewed interests in PR targeting therapies, there is still a lack of unified understanding of the intrinsic activities of PR on whether it is pro-tumoral or anti-tumoral. Two factors are known to impact manifestation of PR activity in response to progestin. The first one is the duration of treatment and the second is the expression levels of PR. It is a well-known paradigm that progestin activated PR elicits biphasic effect on cell proliferation, in which progestin promotes first cell cycle after treatment but induces G0/G1 arrest subsequently in T47D cells and MCF-7PR cells with PR transfection [17-19]. Both T47D cells and MCF-7PR cells express high levels of PR, and the long-term treatment of progestin of these two cell lines induces sustained growth inhibition. It was also reported that progestin induced growth arrest in the ER and PR negative cells MDA-MB-231expressing high levels of PR [20, 21]. This is contrast to the parental MCF-7 cells, in which PR expression is estrogen dependent and PR levels would be very low in hormone-deprived medium. In this scenario, progestin’s effect could only be demonstrated at high concentrations, which is prone to crosstalk with AR, GR and even ER, resulting in growth stimulatory effect [22, 23]. In the presence of estrogen, progestin inhibits estrogen stimulated growth [15, 19]. These studies suggest that the long-term effect of progestin is growth inhibitory in cells with high PR expression.

The initial stimulation of cell cycle seems to be resulted from progestin induced cytoplasmic crosstalk of PR with c-Src-ERK signalling. The proline rich region in the N terminal domain of PR was found to interact with SH3 domain of c-Src and this activated c-Src-ERK signalling for cell proliferation [24, 25]. Whilst the mutation in DNA binding domain retains the activity of PR on thymidine incorporation, PR mutation in proline rich region abolished the effect [26, 27]. Notably, a recent study indicates that PR recruitment in response to progestin to highly accessible enhancers was dependent on activation of ERK signalling, and the activation of the kinase-dependent highly accessible PR enhancers drives the initial cell cycle progression. [28]. However, molecular mechanisms for progestin induced cell cycle arrest after the initial cell cycle promotion are poorly understood. We reported that prolonged progestin treatment in MCF-7PR cells caused replicative senescence and apoptosis [19]. However, this was associated with significant downregulation of caspase 7, caspase 8 and PARP1, indicating complex activities of agonist activated PR. The overarching goal of this study is to achieve a comprehensive understanding of molecular effects of agonist activated PR in MCF-7PR cells through integrated proteomic and transcriptomic analysis. By systematically examining gene and protein expression networks, this research aims to uncover key and novel molecular pathways influenced by PR activation. A deeper insight into these regulatory mechanisms is crucial for advancing PR-targeted therapies, ultimately improving treatment strategies for breast cancer.

## Results and discussion

### Agonist activated PR is strongly growth inhibitory and proapoptotic in MCF-7PR cells

The aim of study was to obtain a comprehensive profiling of PR mediated molecular effects of pure progestin with prolonged treatment. To enable robust detection of PR mediated molecular changes, PR expression in MCF-7 cells is elevated by transfection and the cell line is designated as MCF-7PR. We reported that pure PR agonist R5020 induced marked growth inhibition and apoptosis in MCF-7PR cells [19]. In the current study, we characterized the effect of R5020 with a 5-day treatment scheme (72 hours + sub-cultured for another 48 hours) that to be aligned with proteomic analysis. The 5-day treatment with subculture is to minimize the secondary effect of overcrowding in control cells which grow much faster than the R5020-treated cells.

Fig. 1A shows that PR is undetectable in vector-transfected MCF-7C cells cultured in estrogen deprived medium. PRB is the major isoform expressed in MCF-7PR cells transfected with PRB cDNA, and PRA is expressed at a lower level from the second ATG of PRB cDNA. Expectedly, R5020 almost eliminated ERα in MCF-7PR cells after 5 days because agonist activated PR is known to downregulate ERα expression[19]. Importantly, this treatment scheme caused significant cell cycle arrest in G0/G1 and G2/M, and reduction in S phase of MCF-7PR cells (Fig. 1B). Vector transfected control cells (MCF-7C) also showed a small but significant G2/M cell arrest in response to R5020 likely because of low levels of endogenous PR. R5020 also caused significant increase in apoptosis in MCF-7PR based on Annexin V staining, but not in MCF-7C cells (Fig. 1Ci, 1Cii). Crystal violet staining after 2 weeks of culture showed that R5020 treated MCF-7PR cells had much fewer cells compared to vehicle treated controls (Fig. 1D), as dead cells were washed off during medium change. However, apoptotic proteins PARP1, caspase 7, caspase 8 and their cleaved forms were markedly down regulated in R5020 treated cells (Fig. 1E).

**Fig. 1.**
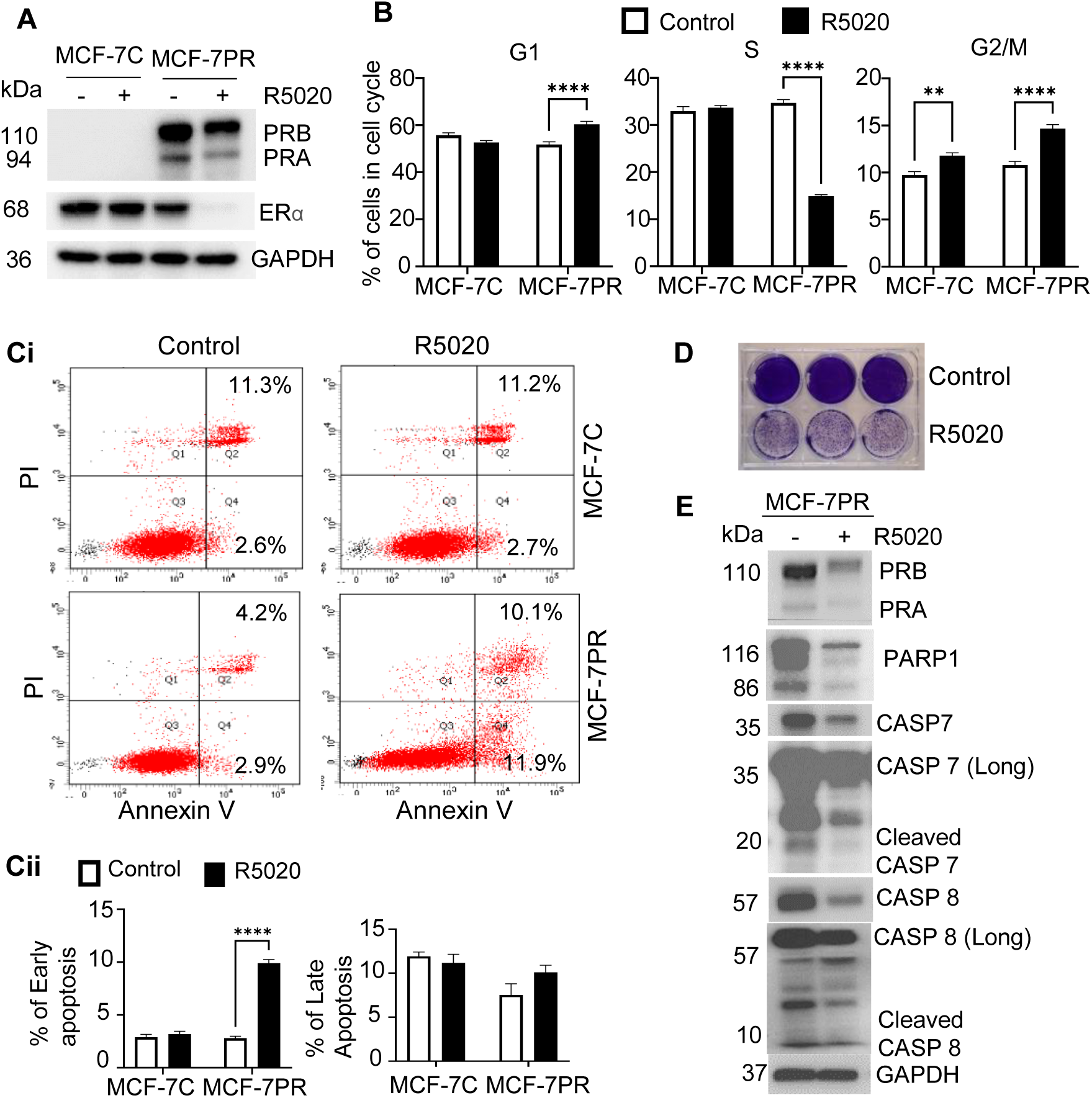
Agonist-activated PR is strongly growth-inhibitory and proapoptotic in MCF-7PR cells. **(A)** Western blot analysis of PR and ERα in vector-transfected MCF-7C cells and PRB cDNA-transfected MCF-7PR cells after 72h + 48h of treatment with R5020. **(B)** Effect of R5020 on cell cycle progression as measured by flow cytometry. Percentages of cells in G1, S, and G2/M phases from two independent experiments (n=6) are presented. **(Ci)** R5020 significantly increased early apoptotic cells (Q4) after 72h +48h of R5020 treatment in MCF-7PR cells. **(Cii)** Percentage of early and late apoptotic cells (n=6 from two independent experiments for MCF-7C cells and n=9 from three independent experiments for MCF-7PR cells). **(D)** Crystal violet staining of cells after two weeks of culture showed that R5020-treated cells had significantly fewer cells compared to controls, as dead cells were washed off during the medium change. (E) R5020 induced downregulation of caspases and PARP1 after 72h +48h of treatment. GAPDH was used as a loading control. All numeric results are presented as mean ± SEM, and the asterisks indicate statistical significance in the comparison between R5020 and the control (* P<0.05, ** P<0.01, **** P<0.0001).

### Agonist activated PR regulates genes in proapoptotic and pro-survival pathways

RNA-Seq analysis was conducted to understand genome-wide effect of R5020 after 72h + 24h treatment of MCF-7PR cells. DESeq2 analysis showed that 6204 genes were significantly regulated (pAdj <0.05) with 3750 genes upregulated and 2454 genes downregulated (Supplementary data 1). The mean average (MA) plot shows generally higher log2 fold changes in upregulated genes than downregulated genes (Fig. 2A). Interestingly, genes coding for basal cell keratins KRT5, KRT6A, KRT6B are among the most significantly upregulated. Known PR target genes such as SGK1, FKBP5, HES2 and HSD11B2 are also among the most significantly upregulated genes.

**Fig. 2.**
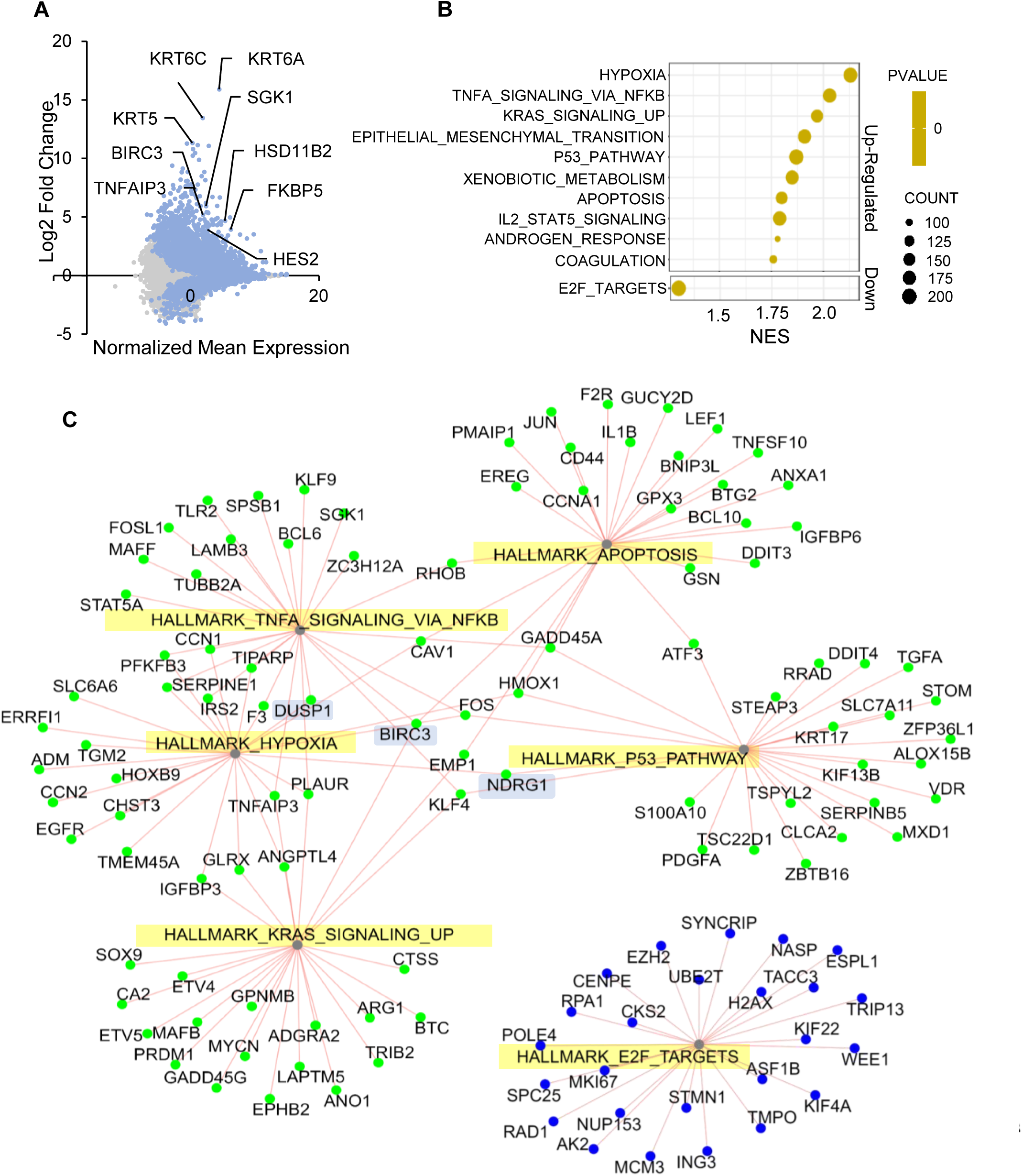
Agonist-activated PR induced enrichment of pro-survival and apoptotic gene hallmarks in RNA-Seq analysis. **(A)** The MA plot shows the normalized mean expression of genes versus the log2 fold change of gene expression between R5020-treated and control samples. **(B)** Bubble plot of GSEA Hallmark pathway analysis, showing the top 10 out of the 32 significantly upregulated hallmarks (p<0.05, FDR<0.05) and one significantly downregulated hallmark (p<0.05, FDR=0.1) by R5020. Due to the limitations of the GSEA software in generating exponential values, the p-value scale shows only one color, as the p-values for all the hallmarks were <0.000. **(C)** Network plot showing the connection of the top 25 genes in the 6 hallmarks, respectively (green node = upregulated, blue node = downregulated).

Gene set enrichment analysis (GSEA) based on Molecular Signatures Database [29, 30] revealed 32 significantly enriched Hallmark gene sets in R5020 treated cells (FDR q<0.05). Top 10 positively enriched Hallmarks (upregulated by R5020) are shown in Fig. 2B. These R5020 induced Hallmarks include those that are pro-survival such as TNFA_SIGNALING_VIA_NFKB and KRAS_SIGNALING_UP, and that are proapoptotic such as HYPOXIA, P53_PATHWAY, and APOPTOSIS. These findings again reflect diverse regulatory functions of PR. Furthermore, some components of these Hallmark gene set also have diverse functions in cancer and can appear in two or more of the Hallmarks as is shown in the network plot of top 25 genes from each of the five Hallmarks (Fig. 2C, Supplementary data 2). The shorter distance between HYPOXIA and TNFA_SIGNALING_VIA_NFKB hallmark indicates that they have the highest number of overlapping genes (10 out of 25). Some of these genes have been reported to have opposing functions on apoptosis or cell proliferation. For example, DUSP1 (Dual specificity phosphatase 1) is reported to inhibit cell proliferation and induce apoptosis in cancer cells [31]. It could also inhibit NF-κB signalling to promote apoptosis [32]. Another example is BIRC3 (Baculoviral IAP repeat-containing protein 3) which is the component of TNFA_SIGNALING_VIA_NFKB, KRAS_SIGNALING_UP and APOPTOSIS. BIRC3 is also known as cellular inhibitor of apoptosis protein 2 (cIAP2) that inhibits apoptosis through the suppression of caspases such as caspase 3, 6 and 8 [33]. On the other hand, BIRC3/cIAP2 also inhibits the survival of non-small cell lung cancer by inhibiting ERK phosphorylation [34]. NDRG1 (N-myc downregulated gene-1), a component of TNFA_SIGNALING_VIA_NFKB, KRAS_SIGNALING_UP, HYPOXIA and P53_PATHWAY Hallmarks, has been reported to promote cancer progression or suppress metastasis depending on post-translational modifications and subcellular localization [35]. NDRG1 is also one of the notable hypoxia inducible proteins that help cells to adapt to hypoxia by degrading sodium-potassium ATPase, thus preserving ATP for cell survival [36].

Consistent with the effect of R5020 in inducing G1 and G2/M cell cycle arrest, R5020 negatively regulates E2F_TARGETS (downregulated by R5020) (FDR = 0.1). 86 of the 200 genes in this Hallmark gene set were downregulated by R5020 (Fig 2B, 2C, Supplementary data 3). The genes critical for S phase entry and mitosis such as TOP2A, PLK4, CDK1, MYC, AURKB, PCNA, SPC24 and SPC25 were significantly downregulated and CDK inhibitor, CDKN2A was upregulated. Taken together, the global gene expression analysis indicates that PR orchestrates complex gene expression programs that may drive cell cycle arrest/cell death as well as survival.

### Quantitative proteomic analysis of MCF-7PR cells reveals highly reliable data sets indicative of global molecular functions of agonist activated PR

Combining transcriptomic and proteomic analysis enables a comprehensive understanding of gene regulation and translational outcome. This integrated approach can also shed light on post-translational modifications and regulatory mechanisms. Tandem Mass Tag (TMT) proteomics was therefore conducted to profile PR regulated proteins and phosphorylation. Triplicates dishes of MCF-7PR cells were treated with or without 10 nM R5020 for 72+48h. Total protein lysates (100 μg) were subjected to in-solution digestion before labelling with the TMT-6plex Isobaric Label Reagents and analysis by LC-MS/MS. The PCA plot (Fig. 3A, Supplementary data 4) shows that the triplicate data within control (C1, C2, C3) and R5020-treated (T1, T2, T3) samples are tightly clustered together, indicating the consistency of the data. The strong effect of R5020 on protein changes are indicated by distinctly clustered controls and R5020 treated samples. A total of 4915 proteins were found to be significantly regulated with 2168 proteins up and 2747 proteins downregulated (p<0.05) (Supplementary data 5). 416 proteins showed significant changes in phosphorylation that involved 678 phosphorylated peptides, of which 275 upregulated and 403 downregulated (Supplementary data 6). Notably, R5020 induced protein changes are largely concordant with mRNA changes. About 60% of the differentially regulated proteins were also correspondingly regulated at the mRNA level (Fig. 3B). Regression analysis of Log2 fold changes of mRNA and protein showed that the two sets of data are significantly correlated (R^2^ =0.4, p=2.24E-13) (Fig. 3C).

**Fig. 3.**
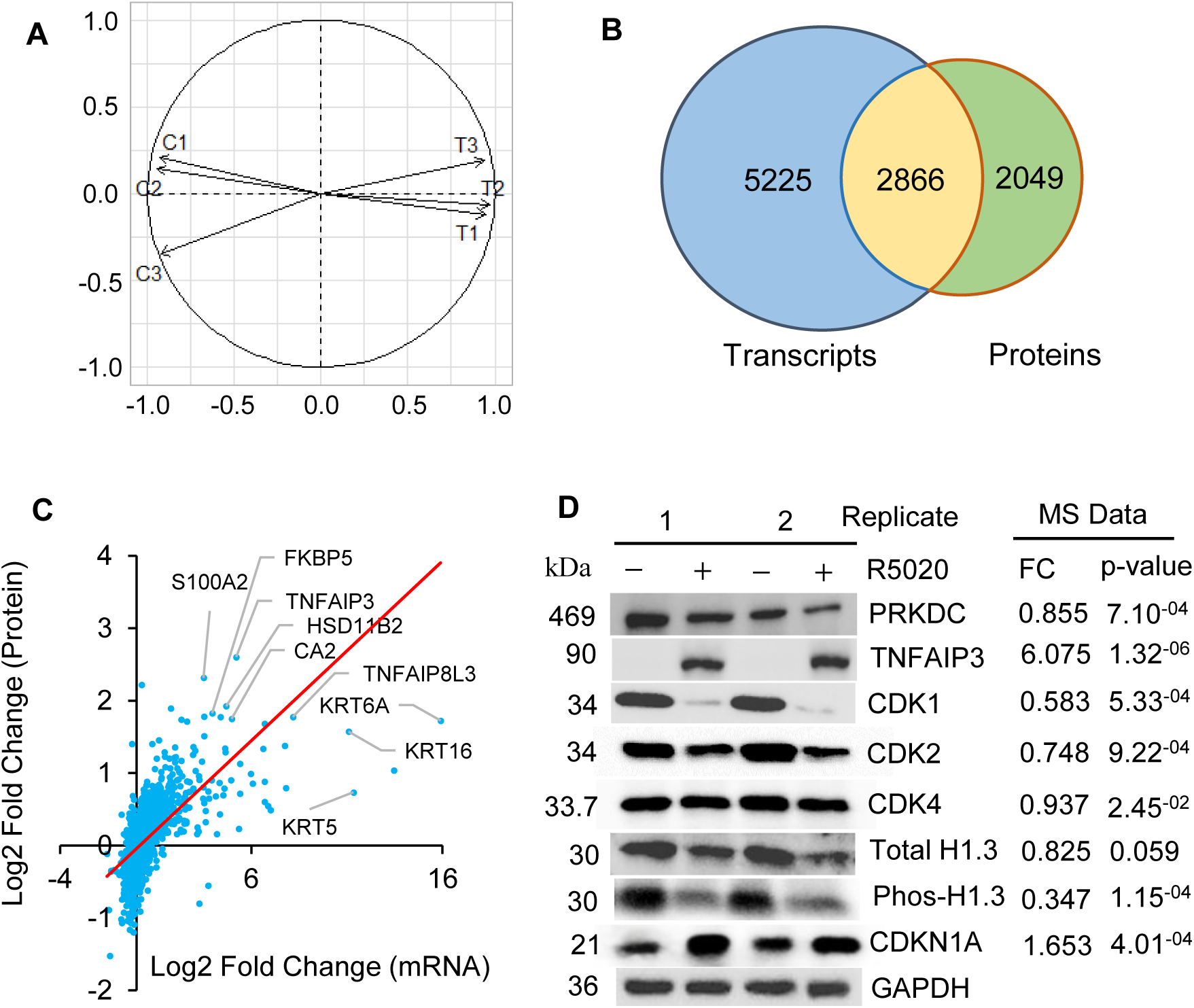
TMT-based proteomics analysis of PR-regulated proteins produced highly reproducible data, indicative of the diverse molecular functions of agonist-activated PR. (A) The PCA plot shows the consistency of the triplicates in the control (C1, C2, C3) and R5020-treated (R1, R2, R3) samples, as evidenced by the distinct clusters. **(B)** The Venn diagram shows the number of differentially expressed genes in transcriptomic (blue) and proteomic (green) data, as well as the overlapping genes/proteins (yellow). **(C)** Regression analysis of transcriptomic and proteomic data shows a significant correlation between mRNA and protein changes. **(D)** Validation of proteomic data by Western blot analysis in replicate samples. The fold change (FC) and p-values of MS data for each protein are shown on the right. GAPDH was used as a loading control for Western blot analysis.

The TMT quantification data are also highly reproducible. Fig 3D shows 100% validation of 7 proteins (DNA-PKcs, TNFAIP3, CDK1, CDK2, CDK4, CDKN1A, H1 histone and its phosphorylated form) by Western blotting analysis. Notably, even a small change in MS quantification (7% reduction in the case of CDK4) is evidently detectable in Western blots. As will also be seen in the subsequent sections of the manuscript, R5020 induced changes in proteomic data can be confirmed in all cases by Western blotting analysis. Thus, our TMT proteomic analysis provides a large valuable dataset showing whole proteomic functions of agonist activated PR in breast cancer cells.

### Agonist activated PR induced broad downregulation of proteins critical for G1-S progression and mitosis

The volcano plot in Fig. 4A highlights top 10 up and downregulated proteins by log2 fold change. All except for KRT71 and CDSN of the top 10 upregulated proteins are also greatly upregulated at the mRNA level. Interestingly, 5 of the top 10 most downregulated proteins are important cell cycle regulators. These are PCLAF (PCNA clamp-associated factor), TIPIN (TIMELESS-interacting protein), TOP2A (Topoisomerase IIα), ISG15 (S-phase kinase associated protein 2) and WEE1. Accordingly, E2F TARGETS, G2/M CHECK POINT and MITOTIC SPINDLE are the top Hallmarks negatively regulated by R5020 (Fig. 4B).

**Fig. 4.**
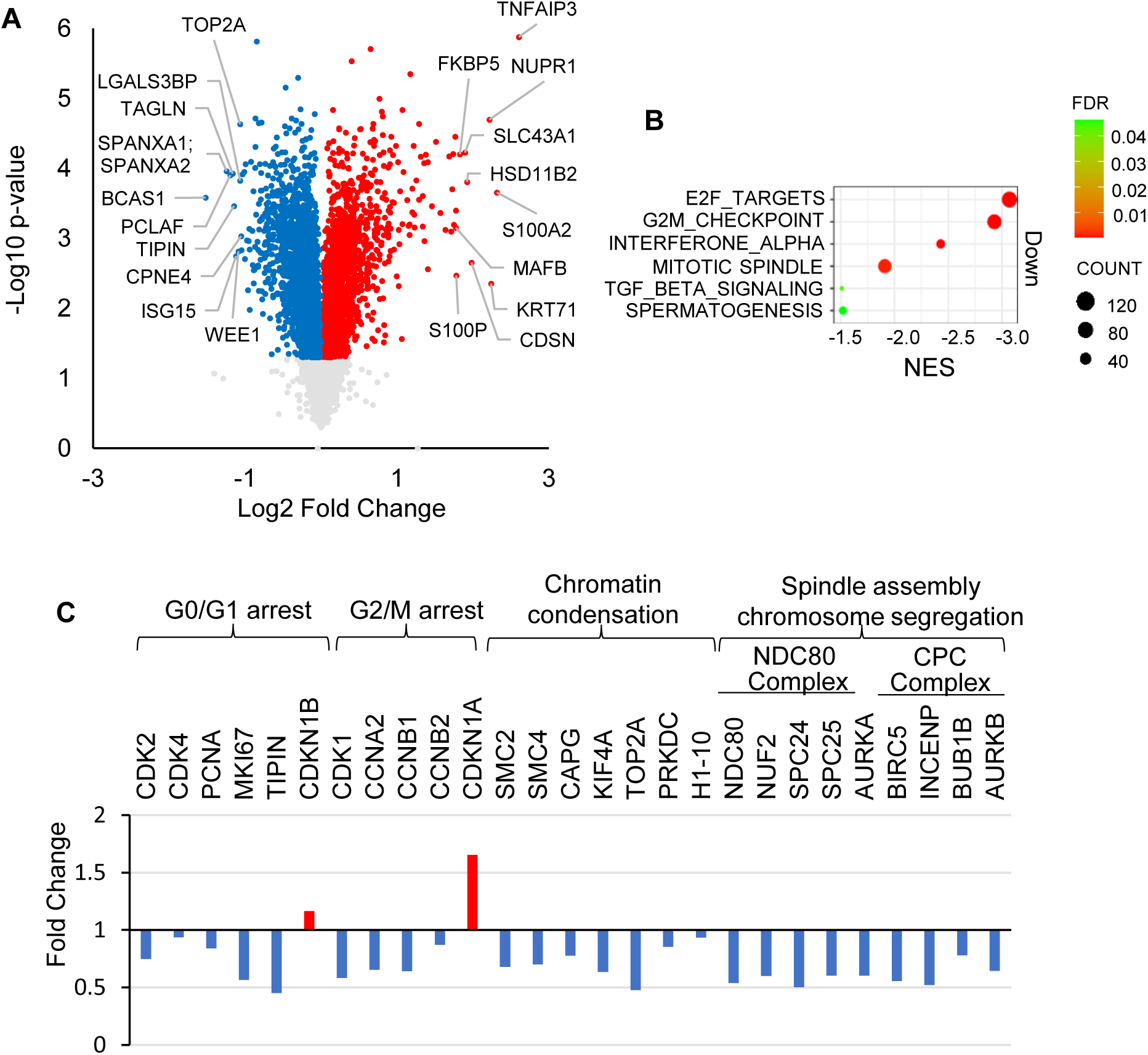
Agonist-activated PR induced broad downregulation of proteins critical for G1-S progression and mitosis. **(A)** Volcano plot of total master proteins with high FDR confidence (Total master proteins = 7783; Significantly regulated proteins = 4915; Significantly upregulated = 2168; Significantly downregulated = 2747). The top 10 up- and downregulated proteins based on fold change are labeled. **(B)** GSEA revealed 6 negatively regulated (FDR < 0.05) Hallmark protein sets by R5020. **(C)** Significantly regulated cell cycle regulators by R5020 in proteomics data. Downregulation of positive regulators is shown in blue bars; upregulation of negative regulators is shown in red bars.

A closer examination of downregulated proteins in these Hallmarks shed light on mechanisms for R5020 induced G0/G1 and G2/M phase arrest (Fig. 4C). Decreases of CDK2 and CDK4, along with an increase of the CDK inhibitor CDKN1B (p27) is consistent with R5020 induced G0/G1 cell arrest as these proteins are crucial for the initiation of DNA replication and the G1-S phase transition. These changes were coupled with significant downregulations of MKi-67 (Marker of Proliferation), PCNA (Proliferating cell nuclear antigen), and TIPIN (TIMELESS-interacting protein), which are essential for DNA replication and DNA damage checkpoint responses [37-39]. Downregulation of CCNA2, CDK1 and CCNB1/2, which form the pre-mitosis promoting factor, is concordant with R5020 induced G2/M phase arrest.

Furthermore, MS data show broad downregulation of proteins involved in chromatin condensation, spindle assembly and chromosome segregation in R5020 treated cells (Fig. 4C). First, proteins critical for chromatin condensation including SMC2, SMC4, CAP-G, TOP2A, KIF4A, PRKDC and H1 linker proteins were significantly downregulated. TOP2A resolves DNA supercoils and entanglements, facilitating smooth chromosome folding. KIF4A, a motor protein organizes chromosome axes together with condensin I to ensure proper compaction [40]. Histone H1 binds to linker DNA and promotes higher-order chromatin folding, whereas PRKDC (DNA-dependent protein kinase catalytic subunit) regulates DNA repair and chromatin remodelling, ensuring chromosomal integrity. Second, all components of Ndc80 complex, Ndc80, Nuf2, Spc24, and Spc25 were significantly downregulated by R5020. The Ndc80 complex is a key kinetochore binding component that links mitotic spindle with centromere-associated proteins for spindle assembly and chromosome alignment [41]. Its downregulation leads to failure of chromosome alignment and segregation during mitosis. Third, all components of Chromosomal passenger complex (CPC), Servivin (BIRC5), inner centromere protein (INCENP), BUBR1 (BUB1B), AURKA and Aurora B (AURKB) were significantly downregulated. These proteins function together to enable phosphorylation of a number of proteins to ensure proper spindle assembly and chromosome segregation [42]. Together, R5020 induced cell cycle arrest at G0/G1 phase and mitosis is mediated by downregulation of CDKs, cyclins and proteins required for DNA replication, chromatin condensation, spindle assembly and chromosome segregation.

### R5020 induced some pro-growth signalling proteins but functionally unproductive

R5020 also induced protein changes that are seemingly growth stimulatory. For example, WEE1 is one of the top 10 most downregulated proteins by R5020 (Fig. 5A). This downregulation in theory promote mitosis because WEE1 prevents the activation of the CDK1-cyclin B complex by inducing inhibitory phosphorylation of CDK1 at tyrosine 15 (Y15) [43]. Indeed, there was about 50% reduction in Y15 phosphorylation of CDK1 in R5020 treated cells (Fig. 5B). However, there is evidence to suggest that reduced WEE1 did not lead to more active CDK1. First, there was more than 50% decrease of CDK1 protein. Hence, the reduced Y15 phosphorylation is likely due to the decrease in CDK1 protein. Second, CDK1 protein activity was markedly reduced as is evidenced by 4-fold reduction (4.02E-04) of NPM1 (Nucleophosmin 1) phosphorylation at Thr199 (Fig. 5C), which is catalysed by CDK1 for the initiation of centrosome duplication [44-46]. Hence, downregulation of WEE1 was not translated into an increased activity of CDK1.

**Fig. 5.**
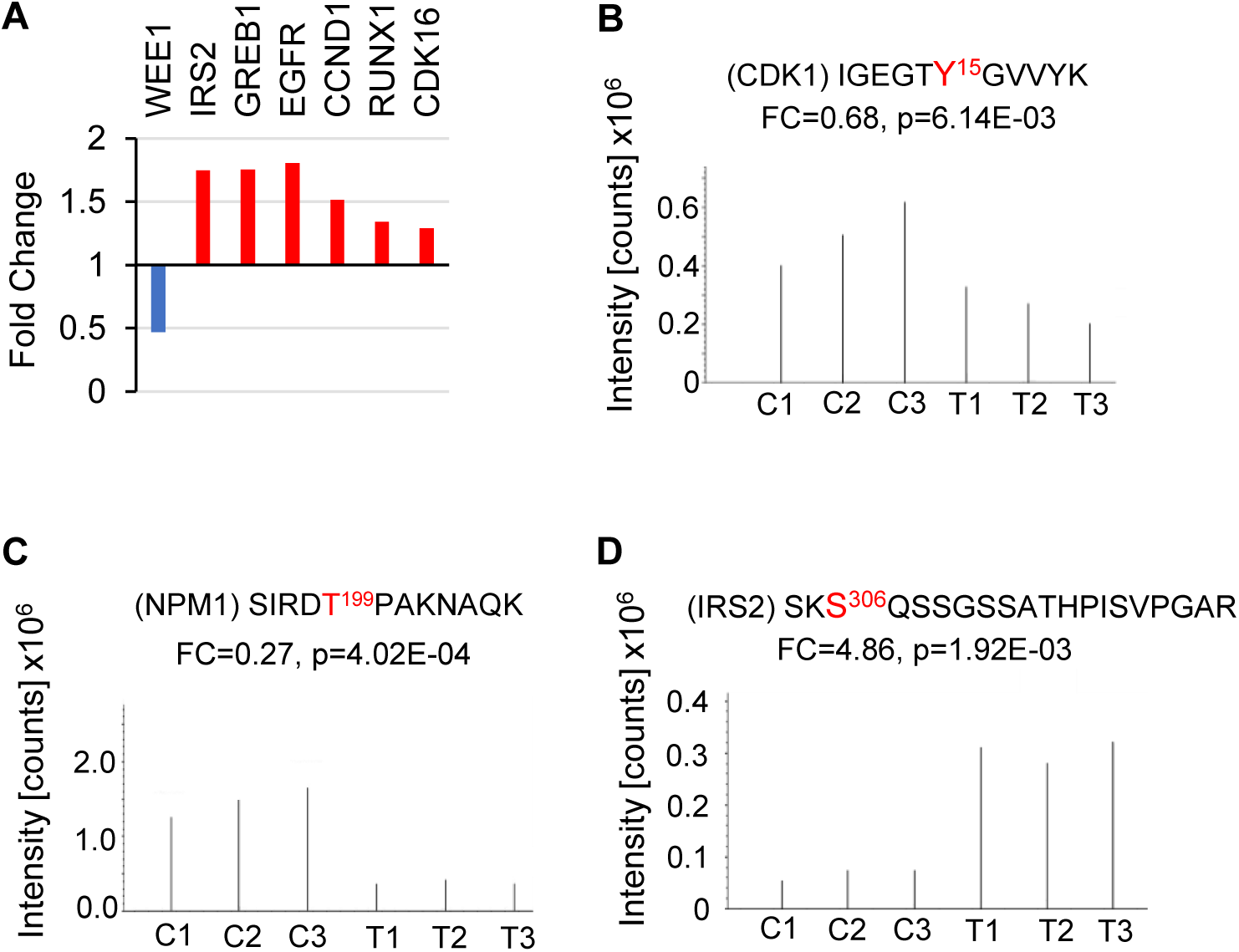
R5020 induced pro-growth signaling proteins. **(A)** The bar plot shows fold changes in protein levels (Blue bar = Downregulation, Red bar = Upregulation). **(B, C, D)** MS spectra of phosphorylated peptides of CDK1, NMP1, and IRS2, respectively. Charts were generated from the raw mass spectrum for better visualization (C = control, T = R5020-treated). The amino acid in red is the site of phosphorylation (FC = Fold Change, p = p-value).

R5020 also induced upregulation of some growth-promoting proteins and protooncogenes such as IRS2, GREB1, EGFR, CCND1, RUNX1 and CDK16, which were also upregulated at the mRNA level (Fig. 5A). IRS2, GREB1, EGFR and CCND1 are reported to be regulated by progestin in breast cancer cells and used as evidence for progestin’s pro-tumoral effect when they were studied individually [47-50]. Our data proves that upregulation of these proteins is in fact functionally inconsequential because R5020 is intensely growth inhibitory. One explanation for the lack of pro-growth function is post-translational modification. For example, although R5020 increased IRS2 protein level by 1.7-fold (Fig. 5A), it also induced 4.86-fold (p=0.00192) increase of the inhibitory phosphorylation of IRS2 at S306 (Fig. 5D), which is phosphorylated by AKT as a negative feedback mechanism [51]. Hence, the upregulation of a pro-growth protein by a PR agonist should not be interpreted as evidence of its pro-growth activity.

### Agonist-activated PR induced enrichment of HYPOXIA, p53 PATHWAY and TNFA_SIGNALING_VIA_NFKB Hallmarks at the protein level

Gene set enrichment analysis (GSEA) of proteomics data revealed 18 Hallmarks protein sets enriched (upregulated) in R5020 treated cells (FDR<0.05). Top 10 are shown in Fig. 6A. HYPOXIA, P53_PATHWAY and TNFA_SIGNALING_VIA_NFKB are top 3 Hallmark protein sets enriched in R5020 treated cells. These are also the top Hallmark gene sets regulated at the mRNA levels. About 73%, 84% and 70% of the proteins in HYPOXIA, P53_PATHWAY and TNFA_SIGNALING_VIA_NFKB Hallmarks overlap with the genes in RNA-Seq Hallmarks (Fig. 6B), indicating that progestin regulated these Hallmarks at the transcription level. Cellular hypoxia is conventionally characterized by insufficient oxygen supply and primarily mediated by hypoxia inducible factors (HIFs) that regulate genes involved in angiogenesis, metabolic adaptation and cell survival [52, 53]. Protein network with top 25 most regulated proteins shows that HYPOXIA share common components with P53_PATHWAY and TNFA_SIGNALING_VIA_NFKB (Fig. 6C). This is not surprising as all three pathways can be activated in response to hypoxic conditions. Hypoxia inducible factor 1A (HIF1A) can also physically interact with p53 and NFKB to regulate their stability under hypoxia or cellular stress. It is conceivable that PR plays an important role in coordinating the activities of these three pathways. Data in the subsequent sections will explain how each of these three pathways may be involved in PR agonist induced apoptosis independent of caspases.

**Fig. 6.**
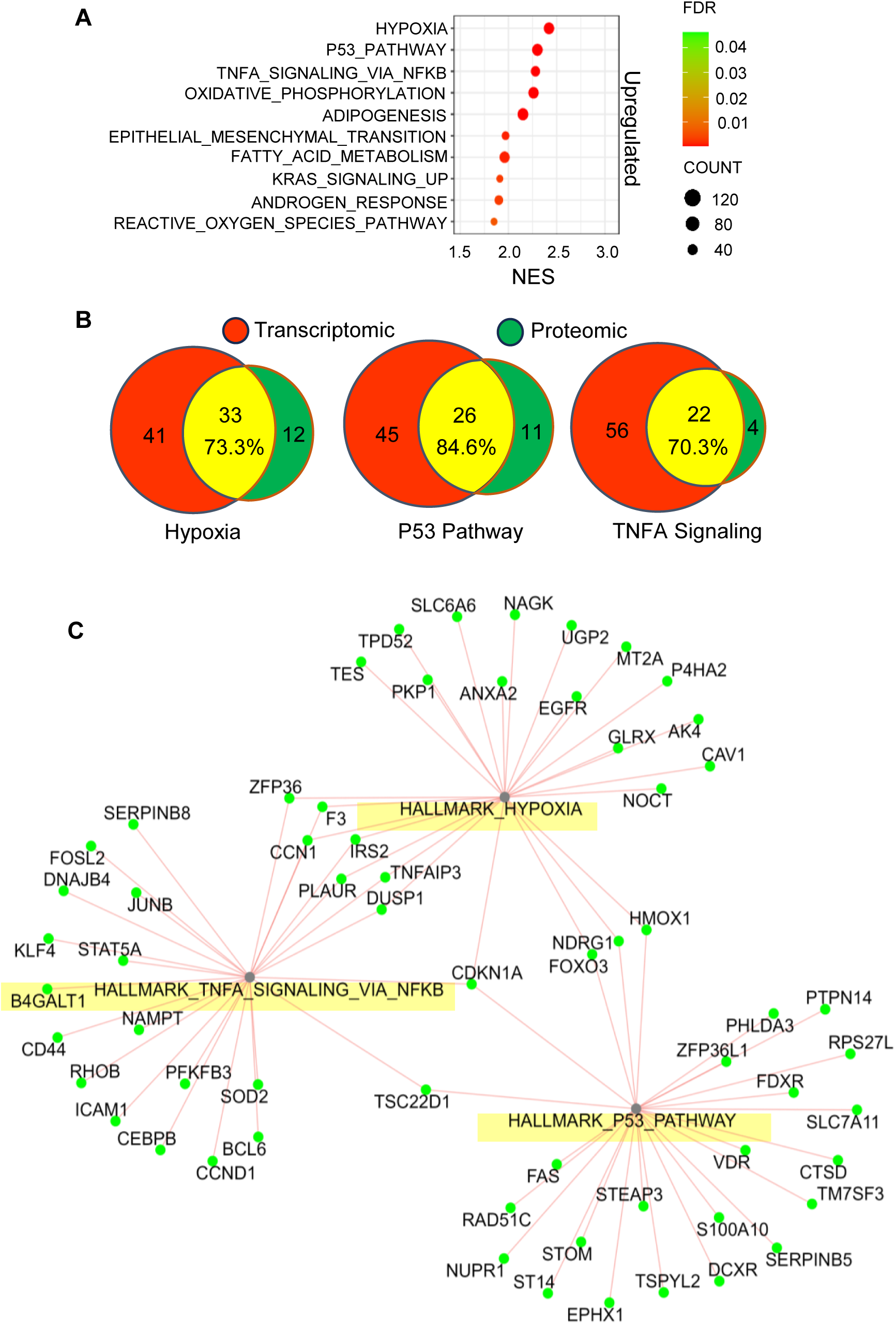
Agonist-activated PR induced enrichment of protein hallmarks with diverse functions. **(A)** GSEA of protein changes revealed 18 significantly upregulated hallmark gene sets by R5020 (p<0.05, FDR<0.05), with the top 10 shown in the bubble plot. **(B)** The Venn diagram shows that the top 3 PR-regulated mRNA and protein hallmarks are highly correlated. The yellow region shows the number and percentage of overlapping proteins with the genes from RNA-seq data. **(C)** The network plot shows the connection of the top 25 proteins in the top 3 upregulated hallmarks, respectively (green node = upregulated).

### R5020 induced the activation of HIF1A–BNIP3/NIX axis causes mitochondrial dysfunction and mitochondrial outer membrane permeabilization (MOMP)

#### R5020 induced upregulation of HIF1A, BNIP3 and NIX

Since R5020 induced HYPOXIA Hallmark under normoxia condition, we examined whether R5020 induced HIFs, master regulators of hypoxic response. It was found that HIF1A was only upregulated by R5020 at the mRNA level (log2 fold change=0.69, pAdj=9.69E-44) after 72+24h treatment. Although HIF1A was not detected in MS data, Western blotting analysis showed that it is massively and progressively increased by R5020 after 24h, 48h, 72h and 72h+24h treatment (Fig. 7A). The increase of HIF1A tapered off at 72h+48h and 72h+72h time points. This was associated with a significant increase of EGLN1 (prolyl hydroxylase domain 2) protein from MS data (1.27-fold, p=0.007). Since EGLN1 hydroxylatesHIF1A for degradation by VHL, HIF1A reduction at later time points is conceivably caused by increased hydroxylation by EGLN1 for degradation. Thus, HIF1A is likely the master mediator of R5020 induced HYPOXIA.

**Fig. 7.**
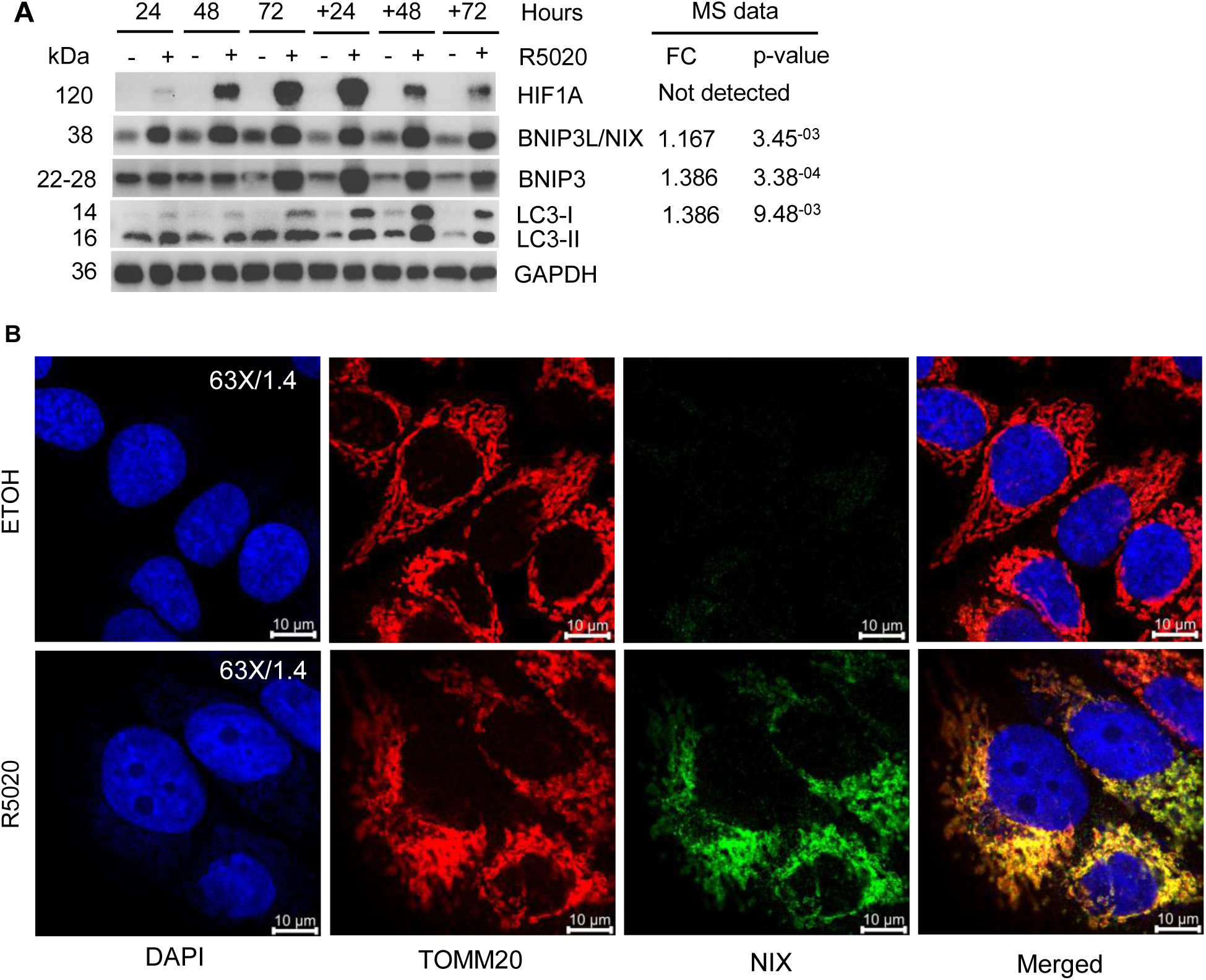
R5020 activates the HIF1A-BNIP/BNIP3L axis. **(A)** Western blot analysis showed that R5020 upregulates HIF1A and its downstream effectors, BNIP3L/NIX, BNIP3, LC3-I, and LC3-II in a time-dependent manner. GAPDH was used as a loading control. **(B)** Co-immunostaining of DAPI, TOMM20, and NIX at 72h of treatment. NIX (green) was detectable and co-localized with TOMM20 (a mitochondria marker, red) in R5020-treated cells, compared to control cells, in which NIX was undetectable.

R5020 also induced the upregulation of the mRNA of BNIP3 (Bcl-2/adenovirus E1B 19 kDa interacting protein 3) (log2 fold change=0.53, pAdj=3.2E-29) and NIX/BNIP3L (will be referred as NIX) (log2 fold change=1.43, pAdj=4.3E-225). BNIP3 and NIX were also upregulated in MS data (BNIP3, Fold change=1.39, p=0.0003; NIX, Fold change=1.17, p=0.003). On Western blot (Fig. 7A), both BNIP3 and NIX were consistently and massively upregulated from day 1 (24h) to 6 (72h+72h). Immunofluorescence shows that NIX/BNIP3L staining colocalizes with mitochondria marker TOMM20 in R5020 treated cells, but it is undetectable in untreated controls (Fig. 7B).

BNIP3 and NIX are known HIF1A target genes. It is logical to assume that they are induced by HIF1A. However, both BNIP3 and NIX were significantly upregulated by R5020 at the mRNA level after 6 hours (data not shown) when HIF1A was not yet significantly upregulated. It is likely that BNIP3 and NIX are also direct target genes of PR. Both PR and HIFA were involved in the progressive upregulation of these two proteins.

#### Agonist-activated PR induced mitochondrial fragmentation and loss of mitochondrial membrane potential

BNIP3 and NIX are members of the BH3-only protein family that promote autophagy and cell death. These proteins can target damaged mitochondria for degradation by interacting with components of the autophagic machinery such as LC3 (microtubule associated protein 1 light chain 3)[54, 55]. This is consistent with previous report that R5020 induces autophagy in MCF-7PR cells [56]. We also observed upregulations of LC3-I and LC3-II at various time points in association with increases of BNIP3 and NIX (Fig. 7A).

High levels of BNIP3 and NIX form pores on the mitochondrial outer membrane, where it contributes to the loss of mitochondrial membrane potential and mitochondrial dysfunction [57, 58]. Indeed, JC-1 staining, which measures mitochondrial membrane potential, showed a time-dependent loss of mitochondrial membrane potential (MtMP) in response to R5020 (Fig. 8Ai). About 30% and 48% of R5020 treated cells displayed a loss of MtMP after 72h+48h and 72h+72h treatment, respectively, compared to 8% in control cells (Fig. 8Aii). This was coupled with marked change in mitochondria morphology. At 20X magnification, Mitotracker staining in almost all R5020 treated cells was more dense, more aggregated and polarized compared to the controls (Fig. 8B). At 63X/1.4 magnification, mitochondria in R5020 treated cells are fragmented with highly dense and spherical aggregates, in contrast to the healthy elongated and interconnected networks in control cells. Hence, R5020 induced mitochondria fragmentation and loss of membrane potential likely through upregulation of BNIP3 and NIX.

**Fig. 8.**
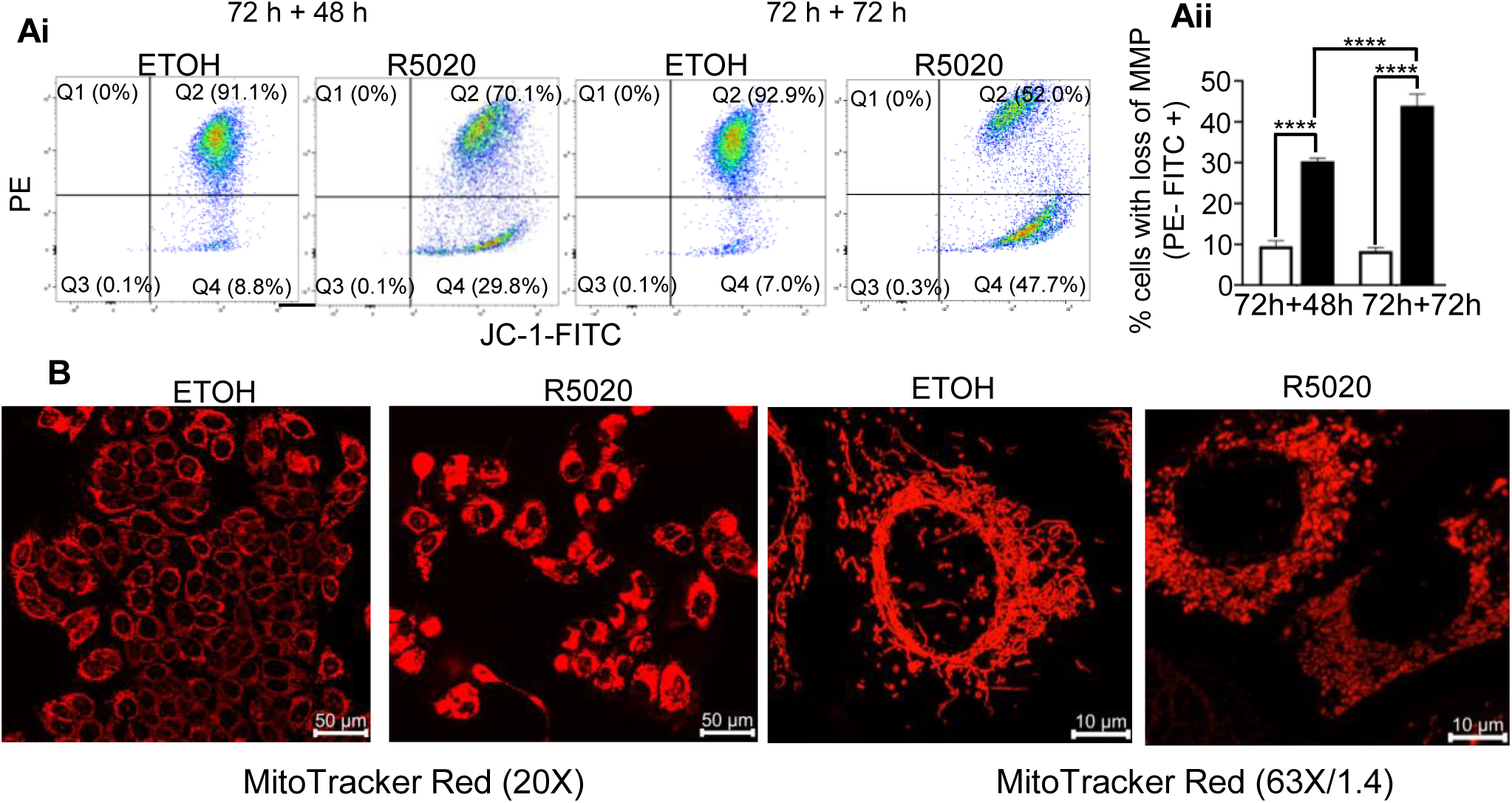
Agonist-activated PR induces mitochondrial fragmentation and loss of mitochondrial membrane potential. **(Ai)** FACS gating strategy for mitochondrial membrane potential using JC-1 staining. Healthy cells show a double-positive signal (Q2 = PE+FITC+), while cells with a loss of membrane potential show a positive FITC signal only (Q4 = PE-FITC+). **(Aii)** Statistical analysis from two independent experiments (n = 6/treatment) showed that R5020 significantly increased the percentage of cells in Q4 compared to the control at both time points. The percentage of cells in the R5020 samples progressively increased across the treatment time points. **(B)** MitoTracker Red was used to observe changes in mitochondrial morphology after 72h + 72h of treatment. R5020 treatment caused the mitochondria to become dense and fragmented into spherical aggregates, in contrast to the elongated and interconnected network in control cells. All numeric results are plotted as mean ± SEM, and the asterisk indicates the statistical significance of the comparison between R5020 and the control (**** P < 0.0001).

#### Agonist-activated PR induced cytochrome c release

Increase of BNIP3 and NIX can contribute to apoptosis through sequestering anti-apoptotic proteins such as BCL-2 and BCL-XL, which normally sequester BH3-only proteins such as BAD, BID, BIM. BH3-only proteins are important for pore formation of BAK and BAX on the mitochondrial outer membrane, leading to mitochondria outer membrane permeabilization (MOMP) [59]. Interestingly, R5020 also significantly reduced the levels of BCL-2 and BCL-XL (Fig. 9A), limiting its inhibition of BH3-only proteins, facilitating pore formation of BAK and BAX. Inactive BAX is localized in the cytoplasm in healthy cells but translocated to the mitochondria in response to apoptotic stress [60]. The composite image at 20X magnification shows that BAX (Green) staining overlaps with TOMM20 (Red) in about 10-15% of cells after 72h+72h R5020 treatment. In contrast, BAX staining is not detectable in control cells which only shows red TOMM20 staining in the composite image (Fig. 9B). At 63X magnification, BAX appears as clustered punctate. Some BAX overlaps perfectly with TOMM20 as yellow punctate, whereas other BAX partially overlap or are adjacent to mitochondria. These clustered BAX punctate are believed to be at different stages of pore formation on the mitochondria membrane [60].

**Fig. 9.**
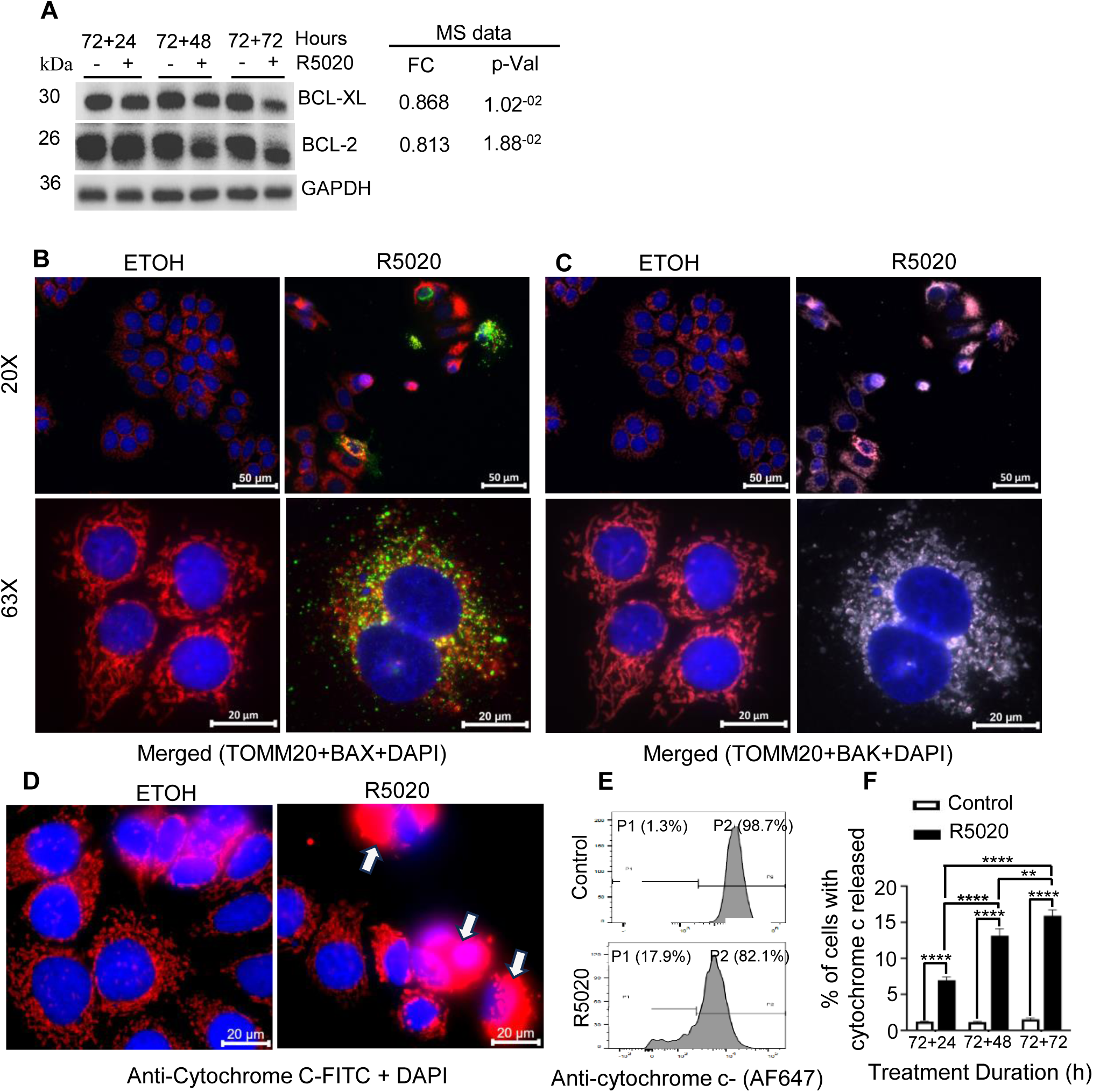
Agonist-activated PR induces cytochrome c release. **(A)** R5020 reduced the levels of anti-apoptotic proteins BCL-XL and BCL-2. GAPDH was used as a loading control. **(B)** Co-immunostaining of TOMM20, BAX, and DAPI at 20X and 63X magnification. In control cells, BAX was undetectable, and only TOMM20 staining (red) was seen in the composite image. BAX (green) appeared as punctate, either fully, partially, or adjacent to the TOMM20 staining (red) in R5020-treated cells. **(C)** Co-immunostaining of TOMM20, BAK, and DAPI at 20X and 63X magnification. R5020 increased BAK punctate (light blue), which largely colocalized with TOMM20 in the composite image (light pink). **(D)** Immunostaining of cytochrome c. The protein was localized to the mitochondria in control cells but was released into the cytoplasm in R5020-treated cells, as indicated by white arrows. **(E)** Gating in FACS analysis of cytochrome c release (P1 = Cells that released cytochrome c; P2 = Cells with cytochrome c in mitochondria). **(F)** Quantification of FACS analysis of cytochrome c. The percentage of cells that released cytochrome c was significantly elevated over time in R5020-treated cells (results from 3 independent experiments, n = 9). All numeric results are plotted as mean ± SEM, and the asterisk indicates the statistical significance of the comparison between R5020 and the control (** p < 0.01, **** p < 0.0001).

On the other hand, BAK mostly colocalizes with TOMM20 in all R5020 treated cells with numerous punctate, whereas control cells only show red staining with TOMM20 because BAK is undetectable (Fig. 9C). However, BAX and BAK punctate partially overlapped (Supplementary Fig 1). This is consistent with the understanding that BAX and BAK can form pores independently although they can also form hetero-oligomeric complexes to create pores in the mitochondrial membrane. In BAX or BAK knockout cells, the remaining protein can form rings independently [61]. Thus, R5020 induces BAX pores in a fraction of cells where BAK punctate were present in almost all cells undergoing mitochondria fragmentation and morphological changes. Non-composite images of BAX and BAK staining with individual antibodies can be found in Supplementary Fig 2 and 3, respectively.

Bax pores in the mitochondrial membrane allow cytochrome c release into the cytosol. Immunofluorescence staining revealed that cytochrome c is evenly distributed in the mitochondria of control cells (Fig. 9D). In contrast, some of the R5020 treated cells have cytochrome c (white arrow) released into cytoplasm showing whole cytoplasmic staining. FACS analysis shows only 1-2% of control cells with cytochrome c release. In contrast, 6.9%, 13.1% and 15.9% R5020 treated cells released cytochrome c after 96h, 120h and 144h treatment, respectively (Fig. 9E, 9F). Note that the percentage of cells with cytochrome c release at 144h was consistent with the percentage of cells with BAX clustered on the mitochondria in Fig. 9B.

### Agonist-activated PR induces mitochondria-mediated apoptosis independent of effector caspases

The release of cytochrome c from mitochondria is a pivotal event in the intrinsic pathway of apoptosis. The released cytochrome c binds to APAF1 to form apoptosomes, which recruit the apoptosis initiator caspase-9 via CARD-CARD domain [62]. This facilitates the activation of caspase 9 through autocatalytic cleavage within the apoptosome. Although TMT proteomics showed marginal (p= 0.095) upregulation of CASP9, the increase is clearly detectable by Western blotting analysis for both pro CASP9 and cleaved CASP9 (35/37KD) after 72h+48h and 72h+72h R5020 treatment (Fig. 10A), confirming unhindered cytochrome c signalling to CASP9.

**Fig. 10.**
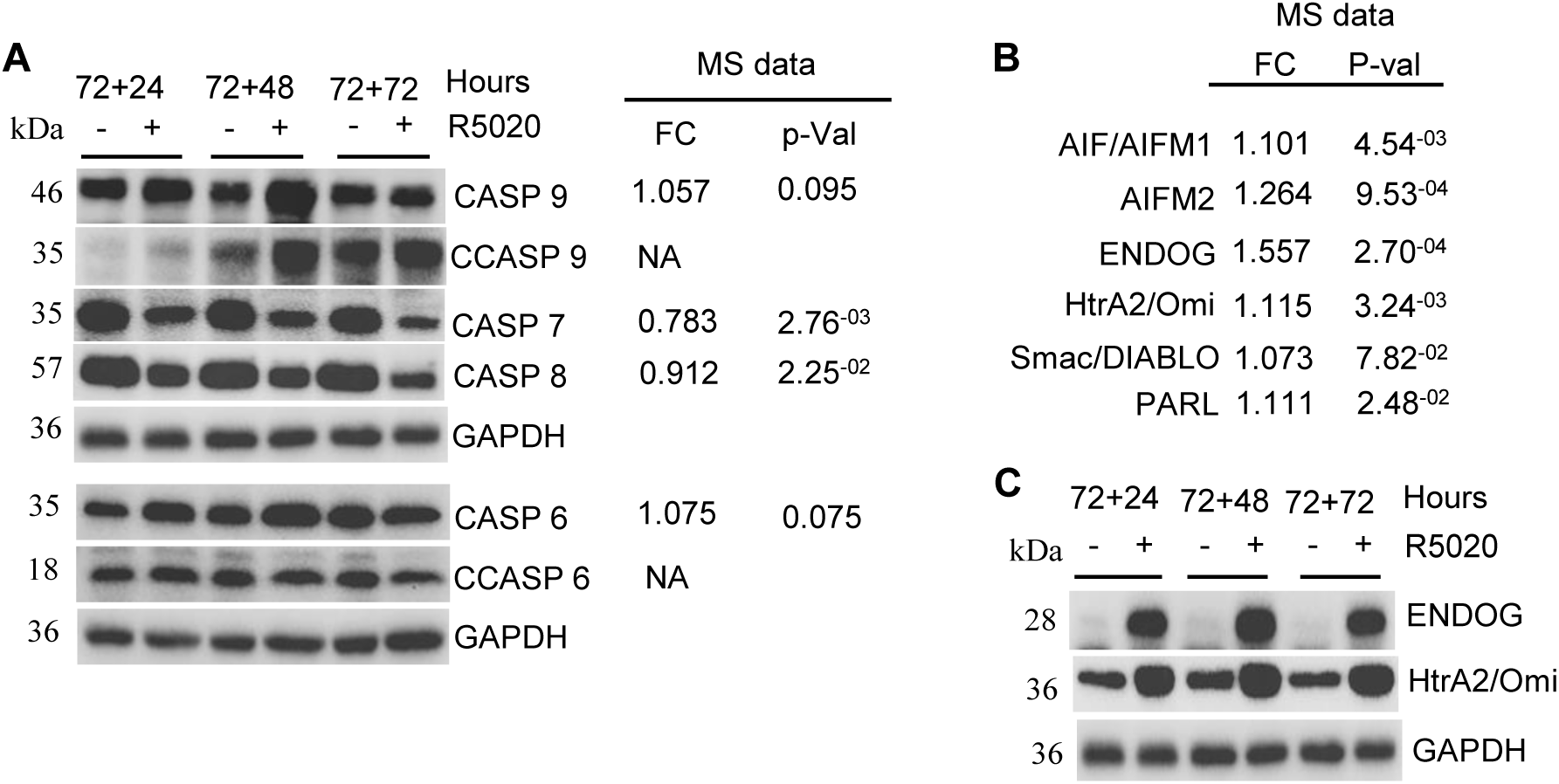
AIF, ENDOG, HtrA2/Omi, and Smac/DIABLO are some of the 200 mitochondrial proteins involved in PR-induced apoptosis independent of caspases. **(A)** Western blot analysis showed the upregulation of the apoptosis initiator caspase 9 but downregulation of the executioner caspases following R5020 treatment. **(B)** Upregulation of mitochondrial pro-apoptotic proteins was found in the TMT proteomics data. **(C)** Two of the mitochondrial pro-apoptotic proteins were validated by Western blot analysis. GAPDH was used as a loading control in all Western blot analyses.

The activated CASP9 is known to directly cleave the effector caspases such as CASP3 and CASP7 for activation [63]. However, the increase of CASP9 was not associated with increases of total or cleaved effector caspases. First, MCF-7 cells are CASP3-defficient [64]. Second, MS data shows that the CASP2 (0.9-fold, p=0.017), CASP7 (0.78-fold, p=0.003) and CASP8 (0.91-fold, p=0.02) were all downregulated, and the decrease of CASP7 and CASP8 were validated again by Western blotting analysis (Fig. 10A). Although CASP6 protein is marginally increased in MS data (1.075-fold, p= 0.07) and validated in Western blots, the cleaved CASP 6 was decreased (Fig. 10A). Thus, R5020 induced increase of CASP9 activation was not associated with increased activity of effector caspases.

Since there is strong evidence for mitochondria mediated cell death but no evidence for increases of effector caspases, we scrutinized MS data for mitochondrial proteins that may be involved in R5020 induced apoptosis. Of 1136 of mitochondrial proteins reported to date [65], 200 mitochondrial proteins were significantly regulated by R5020. 154 proteins were upregulated, and 46 proteins were downregulated (Supplementary data 7, p<0.05). Expectedly, most of the proteins are enzymes involved in metabolism and energy homeostasis. The top 3 upregulated proteins are ACSF2 (Acyl-CoA Synthetase Family Member 2), SOD2 (Superoxide dismutase 2) and AK4 (Adenylate Kinase 4). ACSF2 is involved in mitochondrial fatty acid metabolism. SOD2 converts the harmful free radical superoxide into the less reactive hydrogen peroxide. AK4 regulates the balance of adenine and guanine nucleotide. The top 3 downregulated mitochondrial proteins are DUT (Deoxyuridine 5’-triphosphate nucleotidohydrolase, mitochondrial), MTFR2 (Mitochondrial fission regulator 2), and PDK2 (Pyruvate dehydrogenase (acetyl-transferring)] kinase isozyme 2, mitochondrial).

Further analysis of R5020 upregulated mitochondria proteins show that PR regulated enzymes for ATP production and transport. R5020 increase levels of 6 ATP synthase subunit, 3 COQ (3, 5 and 7) proteins involved in the biosynthesis of coenzyme Q, 8 cytochrome c oxidases (COX) and assembly proteins, 7 mitochondrial ribosomal proteins, 5 subunits of the NADH dehydrogenases (complex I), 9 SLC25A family proteins that are responsible for ATP transport from the mitochondrial matrix, 10 TIMM (translocase of inner mitochondrial membrane) family of proteins and UQCRC1 and UQCRC2 (Complex III). These proteins are all parts of mitochondrial respiratory chain important for oxidative phosphorylation and ATP production.

Importantly, well-documented pro-apoptotic mitochondrial proteins in addition to BCL-2 family of proteins were mostly upregulated. These include Apoptosis-Inducing Factor (AIF/AIFM1), AIFM2, Endonuclease G (ENDOG), High Temperature Requirement A2 (HtrA2/Omi), Second Mitochondria-Derived Activator of Caspases (Smac/DIABLO) and PARL (Presenilin-associated rhomboid-like) (Fig. 10B). Western blotting analysis confirmed that ENDOG and HtrA2/Omi were markedly up regulated by R5020 (Fig. 10C). Since verifications by all Western blotting analysis so far have been successful, we are confident that the upregulation of other mitochondria proteins from TMT proteomic analysis are true.

AIF/AIFM1and AIFM2 belong to the family of apoptosis-Inducing Factors that can localize to the inter membrane space of mitochondria. AIF/AIFM1 and AIFM2, p53 target genes [66], have been reported to induce effector caspase independent apoptosis [67-69]. In response to apoptotic stimuli and MOMP, these proteins are released to cytoplasm through proteolytic processing and translocate to the nucleus where it induces chromatin condensation and DNA fragmentation in a caspase-independent manner. Its effect on DNA fragmentation is shown to be exerted through recruiting DNA nuclease such as ENDOG [70, 71]. Like AIF, ENDOG is localized in mitochondria and released to cytoplasm during MOMP. It is subsequently translocated to the nucleus and catalyses DNA fragmentation in caspase-independent apoptosis [72].

HtrA2/Omi is a serine protease that degrades various substrates in the execution phase of apoptosis [73, 74], it plays a crucial part in the caspase-independent apoptosis pathways. HtrA2/Omi also binds to and neutralizing IAPs, thereby facilitating caspase-dependent apoptosis [75, 76]. Similarly, Smac/DIABLO released from the mitochondria promotes apoptosis by relieving CASP9 from the inhibitory activity of IAPs [77, 78]. The intramitochondrial membrane protease PARL cleaves Smac/DIABLO in response to apoptotic signals, resulting in release and activation of Smac/DIABLO in cytoplasm, which in turn inhibits IAPs and promote apoptosis [79].

Take together, we have provided ample evidence that R5020 induces mitochondria mediated apoptosis through upregulation of proteins such as BNIP3 and NIX to induce loss of mitochondrial potential and MOMP. This is associated with significant alteration of a total of 200 mitochondrial proteins including AIFM1, AIFM2 and ENDOG that are recruited to the nucleus for chromosome condensation and DNA fragmentation. This is also couple with upregulation of proteins such as HTRA2/ OMI, Smac/DIABLO and PARL that inhibit IAPs. Among the many R5020 regulated mitochondria proteins, downregulation of anti-apoptotic proteins BCL-2 and BCL-XL could also play important roles in R5020 induced apoptosis.

### p53 pathway is also involved in PR agonist induced apoptosis

p53 pathway is the second topmost Hallmark gene set enriched n R5020 treated cells. It is well known to induce apoptosis in response to hypoxia, other cellular stress or DNA damage. MCF-7 cells express wild type p53 [80]. We therefore explored whether proteins upregulated in P53_PATHWAY Hallmark mediate R5020 induced apoptosis. Although protein set of P53_PATHWAY is a top Hallmark enriched in R5020 treated cells, p53 protein is not detected in MS data and p53 mRNA did not change either. Western blotting analysis showed that p53 level was downregulated by R5020 treatment (Supplementary Fig 4A). It is possible that the downregulation of p53 results from increased p53 transcriptional activity because DNA binding increases its ubiquitination[81, 82]. P53 is also negatively regulated by the E3 ubiquitin-protein ligase MDM2 which maintain p53 protein at a low level through monoubiquitylation. Although it was not detected in MS data, the expression of MDM2 was upregulated at mRNA level (Log2 fold change=0.38, padj=4.88E-17). This suggests that the feedback mechanism is likely to be the reason why p53 was downregulated even P53_PATHWAY was enriched.

Nuclear Protein 1 (NUPR1), FAS, NDRG1, Stomatin (STOM) and SLC7A11 are the top 5 upregulated proteins in the p53 pathway (Supplementary Fig 4B). NUPR1 is a multifunctional stress-inducible protein that facilitate cell growth and survival. It is reported to protect cells from stress-induced death through RelB and immediate early response 3 (IER3) [83]. It is also a potent ferroptosis repressor through upregulating Lipocalin-2 (LCN2) and gene silencing of LCN2 abolished NUPR1 induced repression of ferroptosis [84]. Although LCN2 protein was not found in MS data, LCN2 was markedly upregulated in gene expression (Log2 fold change=2.02, padj =0.00002) in accordance with upregulation of NUPR1 protein. SLC7A11 is a cystine/glutamate antiporter that also protect cells against ferroptosis by importing cystine into the cell in exchange for glutamate. Cystine is converted to cysteine, a precursor for (GSH) that detoxify reactive oxygen species (ROS) and prevent lipid peroxidation, thereby inhibiting ferroptosis [85, 86]. Therefore, R5020 likely inhibits ferroptosis through regulating NUPR1, LCN2 and SLC7A11.

FAS (CD95) is a cell surface death receptor that is activated by FAS ligand (FASL) to elicit the formation of the death-inducing signalling complex (DISC), leading to the activation of CASP8 [87]. In addition to FAS, R5020 also upregulated death receptors TNFRSF10A (DR4) and TNFRSF10B (DR5) (Supplementary Fig 4B). Although the upregulation of FAS and death receptors was not associated with downstream activation of CASP8, these death receptors may signal to other proteins such as RIP kinases to elicit apoptosis [88].

It is interesting to note that R5020 induced proteins such as AIF/AIFM1 and HTRA2/OMI are direct p53 target genes [66, 89]. Additionally, p53 can translocate to the mitochondria and disrupt the integrity of mitochondria membrane, facilitating the release of SMAC, AIF and ENDOG etc. As mentioned above, these proteins as DNA nucleases and proteases play important roles in apoptosis and DNA fragmentation. Hence, p53 signalling likely contribute to R5020 induced apoptosis.

### PR agonist induced BIRC3/cIAP2 and A20/TNFAIP3 in TNF-**α** signalling via NF-**κ**B did not play a role in downregulation of effector CASPs, PARP1 or apoptosis

We also scrutinized the data to explain the role of TNF-α signalling via NF-κB in apoptosis. NF-κB signalling is a pro survival pathway that activates apoptosis suppressors including inhibitor of apoptosis proteins (IAPs). IAPs are the most important inhibitors of effector caspases [90]. Some IAPs bind to and inhibit caspases, and others such as cIAP1 and cIAP2 are RING finger ubiquitin ligase (E3) that tag caspases for proteasomal degradation [91]. MS data shows R5020 induced downregulation of BIRC2/cIAP1 (0.91-fold change, p=0.009), BIRC5 (Baculoviral IAP Repeat Containing 5) (0.56-fold change, p=1.54E-06) and BIRC6 (0.91-fold change, p=0.012), which is consistent with the pro-apoptotic effect. However, BIRC3/cIAP2 was upregulated by 162-fold at the mRNA level. Although it was not detected by MS, Western blotting analysis showed that BIRC3/cIAP2 protein is hugely upregulated by R5020 (Fig. 11Ai). Since high levels of BIRC3/cIAP2 (referred to as BIRC3 from here) are known to suppress apoptosis through caspase ubiquitination [92-94], we tested whether high levels of BIRC3 is involved in R5020 induced downregulation of CASP7, CASP8 or PARP1. Fig.11B shows that BIRC3 siRNA completely abolished R5020 induced increase of BIRC3, but it had no effect on R5020 induced downregulation of CASP7, CASP8 or PARP1.

**Fig. 11.**
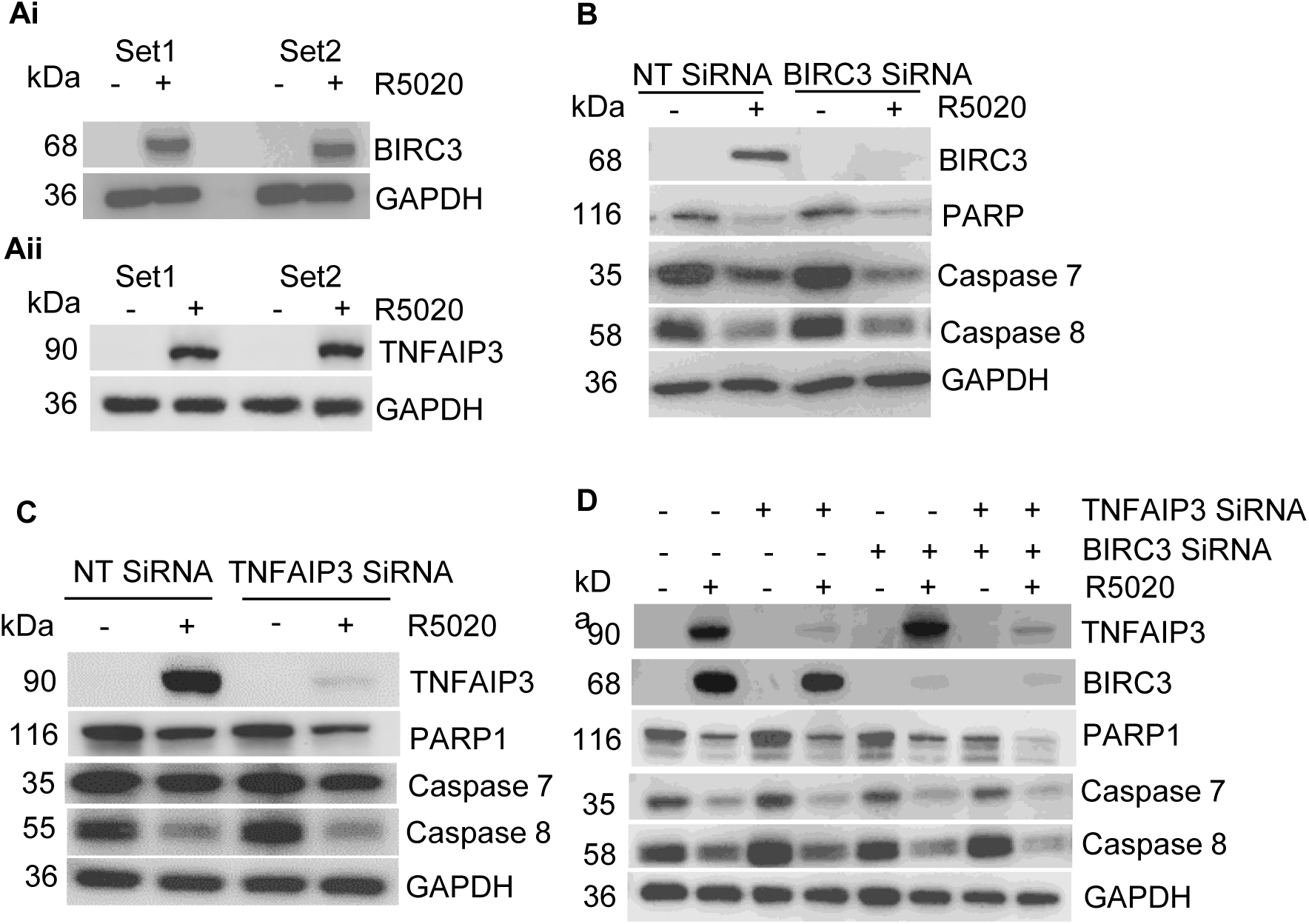
PR agonist-induced BIRC3/cIAP2 and A20/TNFAIP3 did not play a role in the downregulation of CASP7, CASP8, PARP1, or apoptosis. **(Ai)** Upregulation of BIRC3 by R5020 was detected by Western blot analysis, despite not being detected in the MS data. **(Aii)** The R5020-induced increase in TNFAIP3 in MS data was validated by Western blot analysis. **(B, C)** Western blot analysis revealed that BIRC3 and TNFAIP3 siRNA abolished the R5020-induced upregulation of the proteins. However, gene silencing of BIRC3 and TNFAIP3 did not affect the levels of CASP7, CASP8, or PARP1. **(D)** Double knockdown of BIRC3 and TNFAIP3 did not affect the R5020-induced downregulation of CASP7, CASP8, or PARP1 either. GAPDH was used as a loading control in all Western blot analyses.

A20/TNFAIP3 (referred to as TNFAIP3 from here) is another E3 ligase in NF-κB signalling that promotes CASP8 ubiquitination and inhibits apoptosis [95-98]. It is the most upregulated protein by R5020 by fold change and its upregulation is validated in Western blots (Fig. 11Aii). However, TNFAIP3 gene silencing also had no effect on the levels of CASP7, CASP8 or PARP1 (Fig. 11C). Since both BIRC3 and TNFAIP3 are involved putatively in caspase ubiquitination, we asked whether these two proteins could compensate for the loss of effect of the other. Western blots in Fig. 11D shows that BIRC3 and TNFAIP3 were greatly upregulated by R5020. Their respective siRNAs effectively abolished R5020 induced upregulation of these two proteins alone or together. However, the gene silencing of either one or both together did not affect R5020 induced downregulation of CASP7, CASP8 or PARP1 (Fig. 11D). The gene silencing of BIRC3 and TNFAIP3 alone or in combination also did not affect R5020 induced growth arrest or apoptosis (data not shown). Hence R5020 induced BIRC3/cIAP2 or A20/TNFAIP3 is not involved in the regulation of CASP7, CASP8, PARP1, or apoptosis, It is to be notes that CASP7 and PARP1 were significantly downregulated by R5020 at the mRNA level (log2 fold change = -0.32, pAdj = 4.16E-06, log2 fold change = -0.105, pAdj = 0.028, respectively), which suggests their downregulation at the transcription levels may be responsible for their decreases at protein levels.

### Concluding remarks

Current understanding of PR activity in breast cancer cells has largely been derived from studies focusing on individual proteins and pathways. This study employs an integrated omics approach to achieve to date the most comprehensive understanding of PR regulated proteins and molecular networks. Remarkably, the study confidently and reproducibly identified 4,915 PR-regulated proteins and 678 phosphorylated peptides, underscoring PR’s profound and far-reaching influence on various cellular processes. Through detailed mechanistic and phenotypic analyses, the study offers novel insight into proteins and pathways involved in PR driven growth arrest and cell death (Fig. 12). The datasets generated here provide an invaluable resource for developing PR targeted therapies in breast cancer.

**Fig. 12.**
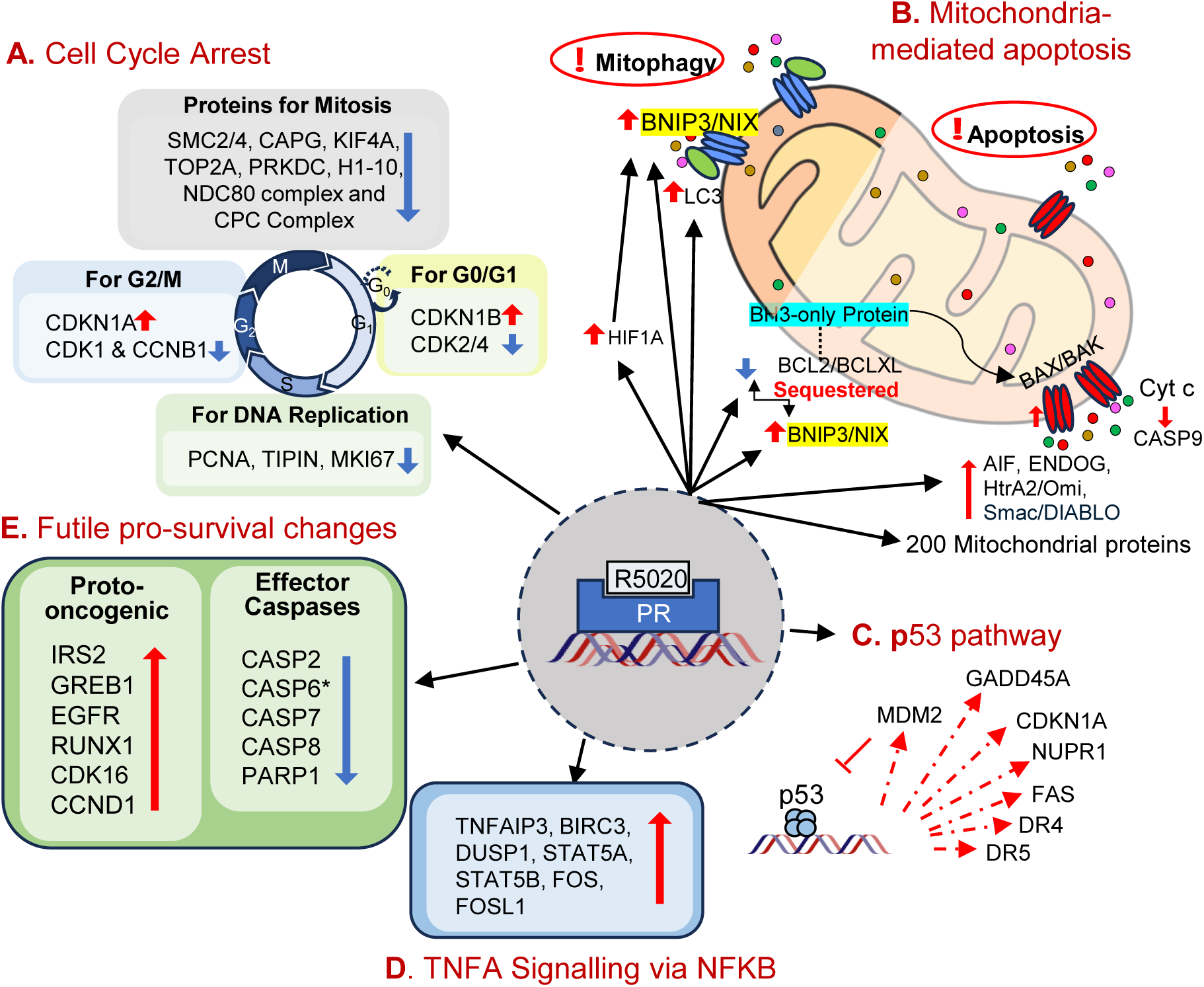
Examples of PR regulated proteins and pathways involved in cell proliferation and apoptosis. The quantitative proteomic analysis identified significant changes of thousands of proteins and many hundreds of phosphorylated peptides in response to PR agonist. This figure shows proteins and pathways that were analyzed and interpreted in greater details. **(A)** Proteins and complexes involved in R5020 induced G0/G1 and G2/M cell cycle arrest, which could also be triggers for apoptosis. **(B)** PR regulated mitochondrial proteins involved in R5020 induced apoptosis. Significant increases of HIF1A and its downstream targets such as BNIP3 and NIX activated mitophagy together with upregulated LC3. Prolonged increase of BNIP3 and NIX causes mitochondria dysfunction and loss of mitochondria membrane potential. BNIP3 and NIX can sequester anti-apoptotic proteins such as BCL-2 and BCL-XL. This relieved BH3-only proteins, which facilitated formation of BAX and BAK pores on mitochondria outer membrane, leading to MOMPs, cytochrome c release and CASP9 activation, along with upregulation and releases of pro-apoptotic factors such as AIF, ENDOG, HtrA2/Omi, and Smac/DIABLO. PR induced apoptotic mechanism may also involve some of the 200 mitochondrial proteins regulated by R5020. **(C)** p53 pathway may also participate in R5020-induced cell apoptosis through upregulation of proteins such as death receptors FAS, DR4 and DR5. Upregulation of MDM2 is involved in a negative feedback regulation of p53. **(D)** TNF signaling via NFKB is generally pro-survival. It was found to be markedly upregulated. Gene silencing of its key players BIRC3 and TNFAIP3 alone or in combination had no influence on R5020 induced downregulation of effector caspases and apoptosis. **(E)** True to the holistic nature of proteomic analysis, the study also detected the previously reported upregulation of protooncogenic proteins by R5020, but the upregulation of these proteins are futile and should not be interpreted as evidence for pro-tumoral activity of PR. Additionally, all effector CASPs detected in the MS data and in immunoblots were downregulation despite R5020 induced apoptosis. *Although CASP6 is marginally increased in MS data and immunoblot, its activation by cleavage is evidently reduced in immunoblot. Blue downward arrow = Downregulation; Red upward arrow = Upregulation.

Importantly, this study revealed for the first time that progesterone receptor (PR) exerts a significant influence on mitochondrial function by regulating a diverse array of mitochondrial proteins involved in various cellular processes. Mitochondria play a central role in critical cellular functions such as apoptosis, metabolism, and the maintenance of redox balance. Given their pivotal role in cell survival and death, mitochondria have emerged as attractive therapeutic targets in cancer research. Recent research has focused on exploiting mitochondrial vulnerabilities in cancer cells to induce selective cell death. Several targeting strategies have been explored, including the induction of MOMP using small-molecule BH3 mimetics, disrupting bioenergetic homeostasis, and targeting the fission and fusion processes to imbalances in mitochondrial dynamics. Remarkably, the study found that agonist-activated PR regulates mitochondrial proteins that are directly implicated in these vulnerabilities. Specifically, PR activation induces MOMP, alters mitochondrial metabolism, and disrupts the balance of mitochondrial fission and fusion dynamics [99-103]. These multifaceted effects suggest that PR activation can interfere multiple mitochondrial pathways. Hence, targeting PR with agonists could offer a “one stone, many birds” approach, achieving multiple therapeutic benefits through a single intervention. Furthermore, the ability of PR to influence mitochondrial function opens new avenues for understanding the broader implications of hormone receptor signalling in cancer biology. Future research could explore the interplay between PR and other mitochondrial regulators, as well as the potential for combining PR agonists with other mitochondrial-targeting agents to achieve synergistic effects. Such combinatorial approaches could pave the way for more effective and personalized cancer therapies

It is also important to note that this study enables a holistic understanding of PR regulated molecular events and offer evidence for reconciliation of current controversies over PR activity in breast cancer. The analysis not only offers functional and mechanistic insights into PR’s antiproliferative and proapoptotic effects but also replicates findings from previous studies reporting proliferative signalling, where PR agonists upregulated oncogenic proteins such as EGFR, IRS2, CCND1, and GREB1 [47-50, 104]. However, upregulation of these protooncogenes and pathways were functionally insignificant, underscoring the importance of cellular context in determining protein activity. These findings reconcile the long-standing controversy surrounding PR function in breast cancer. Furthermore, this study emphasizes the importance of evaluating the effect of drug targets at a global level, particularly in cases where there are conflicting views. Collectively, these findings suggest that breast cancers with high PR expression can benefit from treatment with pure PR agonists. We have tested MCF-7 cells transduced with lower level of PR than MCF-7PR cells and found that the extent of growth inhibition and cell death are related to levels of PR. We suggest that pure PR agonist warrant evaluation as first-line endocrine therapy for breast cancer with high PR expression.

## Materials and methods

### 2.1. Cell Culture

The MCF-7 cells that were transfected with either the control vector pcDNA 3.1 (MCF-7C) or pcDNA 3.1-PRB (MCF-7PR) in a previous study were used throughout this study [19]. The cells were regularly maintained in Dulbecco’s Modified Eagle’s Medium (DMEM) containing phenol red, enriched with 2 mM L-glutamine and 7.5% fetal calf serum (FCS) (Sigma-Aldrich, St. Louis, MO, USA). Cells were incubated in a humidified environment with 5% CO2 and 95% air at 37°C.

### 2.2. Treatment with Promegestone (R5020)

DMEM without phenol red, supplemented with 2 mM L-glutamine and 5% dextran-coated charcoal-treated FCS (DCC-FCS), was used to grow the cells for all experiments involving PR agonist R5020 treatment, except for methods specifically noted. The cells were cultured in this medium for 48 hours before treatment. 100% ethanol (ETOH) was used to reconstitute R5020, which was kept at a stock concentration of 10 mM. The working concentration of R5020 in all experiments was 10 nM, with 0.1% ethanol as the final concentration in the medium. The vehicle control used was 0.1% ethanol. For most experiments, cells were treated with either 0.1% ethanol or 10 nM R5020 for 72 hours. The cells were then sub-cultured in the medium containing the respective treatment and further treated for another 24 hours (72h + 24h), 48 hours (72h + 48h), or 72 hours (72h + 72h), depending on the experimental purpose.

### 2.3. Cell cycle analysis

Cell cycle analysis was conducted to determine the effect of R5020 on cell cycle progression. After the respective treatment conditions, cells were detached using 0.05% trypsin and stained with propidium iodide (PI) in Vindelov’s cocktail [10 mM Tris-HCl (pH 8), 10 mM NaCl, 50 mg PI/L, 10 mg/L RNase A, and 0.1% NP40] for 30 minutes at 4°C. The cells were then analyzed using the BD LSRFortessa™ X-20 flow cytometer (BD Biosciences, Franklin Lakes, NJ, USA) with an excitation wavelength of 488 nm. The data were analyzed with the FlowJo program to obtain the cell cycle distribution of each cell population.

### 2.4. Apoptosis assay

A total of 2 × 10^5^ cells were plated in 60 mm plates for 48h prior to treatment with the 0.1% ETOH or 10 nM R5020. After 72h of treatment, the cells were trypsinized, sub-cultured, and further treated for another 48h. After 72h + 48h of treatment, the cells were collected and stained with anti-Annexin V antibody and propidium iodide (PI) from the Dead Cell Apoptosis Kit with Annexin V Alexa Fluor™ 488 & Propidium Iodide Kit (Invitrogen, Carlsbad, CA, USA), following the manufacturer’s protocol. The stained cells were analyzed using the BD LSRFortessa™ X-20 flow cytometer (BD Biosciences, Franklin Lakes, NJ, USA).

### 2.5. Crystal Violet Staining

The MCF7-PRB cells were first treated with either 0.1% ETOH or 10nM R5020 for 2 weeks, with medium changes every 3 days. A 1% solution of paraformaldehyde (PFA) in phosphate-buffered saline (PBS) was used to fix the cells for 15 minutes, followed by two washes with PBS. The cells were then stained with 0.1% crystal violet in 10% ethanol for 15 minutes at room temperature, followed by several washes with deionized water before air-drying.

### 2.6. RNA-Seq analysis of gene expression

RNA-Seq analysis was performed to determine the effect of R5020 treatment on the global gene expression of the cells compared to the vehicle control after a 96-hour treatment. There were triplicates for each of the treatments (0.1% ethanol or 10 nM R5020). Total RNA was extracted using TRIzol Reagent (Thermo Fisher Scientific, USA) and treated with DNase I using the DNA-free™ DNA Removal Kit (Invitrogen, Carlsbad, CA, USA) to remove any contaminating DNA. The total RNA was sent to the Genome Institute of Singapore, Agency for Science, Technology, and Research, for library preparation and paired-end sequencing.

For quality control, Trim Galore was used to trim the FASTQ files, and FASTQC was performed on these files. STAR was used to map the reads to the human genome GRCh38 using GENCODE 41 annotations. FeatureCounts were then used to generate the gene counts using GENCODE 41 annotations. The gene counts were then used for DESeq2 analysis. Based on the DESeq2 analysis, genes with a p-value less than 0.05 (p < 0.05) were identified as differentially expressed genes (DEGs). Normalized count data from DESeq2 were used for further analysis.

### 2.7. Proteomics

Proteomics was conducted to determine the effect of R5020 on global protein changes and phosphorylation after a 72h + 48h treatment. Each treatment (0.1% ethanol or 10 nM R5020) had triplicates (n = 3) in the proteomics study.

#### 2.7.1. Preparation of proteins (in solution)

A total of 100 μg protein from each condition was subjected to in-solution digestion before labelling the resultant peptides using the TMT-6plex Isobaric Label Reagent Set (Thermo Scientific, Rockford, IL, USA) according to the manufacturer’s protocol. The labelled samples were combined prior to fractionation on a Xbridge™ C18 column (4.6LJ×LJ250LJmm, Waters, Milford, MA, USA) and subsequent analysis by LC-MS/MS.

#### 2.7.2 Liquid chromatography with tandem mass spectroscopy (LC-MS-MS)

The peptides were separated and analyzed using a Vanquish Neo UHPLC System coupled to an Orbitrap Exploris 480 (Thermo Fisher Scientific, MA, USA). Separation was performed on an EASY-Spray 75LJμmLJ×LJ15LJcm column packed with PepMap Neo C18 2 μm, 100 Å (Thermo Fisher Scientific) using solvent A (0.1% formic acid) and solvent B (0.1% formic acid in 80% ACN) at flow rate of 300LJnL/min with a 60 min gradient. Peptides were then analyzed on a Orbitrap Exploris 480 apparatus with an EASY nanospray source (Thermo Fisher Scientific) at an electrospray potential of 2.0LJkV. A full MS scan (350–1,600LJm/z range) was acquired at a resolution of 120,000. Dynamic exclusion was set as 25 s. The resolution of the higher energy collisional dissociation (HCD) spectra was set to 300,00. The automatic gain control (AGC) settings of the full MS scan and the MS2 scan were 300% normalized AGC target and standard, respectively. The Data dependent mode is cycle time. The time between each master scan is 2 seconds. An isolation window of 0.7 m/z was used for MS2. Single and unassigned charged ions were excluded from MS/MS. For HCD, the normalized collision energy was set to 36%.

Raw data files from the three technical replicates were processed and searched using Proteome Discoverer 2.1 (Thermo Fisher Scientific). The raw LC-MS/MS data files were loaded into Spectrum Files (default parameters set in Spectrum Selector) and TMT 6-plex was selected for the Reporter Ion Quantifier. The SEQUEST algorithm was then used for data searching to identify proteins using the following parameters; missed cleavage of two; dynamic modifications were TMT-6plex (+229.163LJDa) (H, S, T), oxidation (+15.995LJDa) (M),, deamidation (+0.984LJDa) (N), Methylation (+14.0156 Da)(R), (+28.03130 Da)(R) and Phosphorylation (+79.966LJDa) (S, T, Y). The static modifications were TMT-6plex (+229.163LJDa) (any N-terminus and K) and Carbamidomethyl (+57LJDa) (C). Percolator is applied to filter out the false MS2 assignments at a strict false discovery rate of 1 % and relaxed false discovery rate of 5 %. The Normalization mode was set based on the total peptide amount.

#### 2.7.3 Proteomics Data Filtering

A total of 8,818 proteins were found in the proteomics raw data before any filtering. Out of these, 8,344 proteins had a high false discovery rate (FDR), and 561 proteins were further removed due to the absence of abundance data. Thus, a list of 7,783 proteins remained for analysis. A p-value of less than 0.05 (p < 0.05) was considered significant for differences in protein expression between ETOH and R5020. An overview of the data was illustrated in a volcano plot, generated using Microsoft Excel.

### 2.8. Gene set enrichment analysis (GSEA)

GSEA analysis was conducted using the software downloaded from its official website https://www.gsea-msigdb.org/gsea/index.jsp. The analysis was performed with the default parameters. An enriched gene set was considered significant if the normalized p-value was less than 0.05. A bubble plot was generated using the online platform SRplot for the GSEA output.

### 2.9. TNFAIP3 and BIRC3 Small Interfering RNA (SiRNA) Transfection

MCF-7PR cells were seeded at a density of 2 ×10^5^ in a 6-well microtiter plate and starved for 24h in phenol red-free medium supplemented with 5% DCC-FCS and 2 mM L-glutamine. The cells were then treated with Non-Targeting Pool siRNA, hTNFAIP3 SMARTpool siRNA, or BIRC3 SMARTpool siRNA by mixing the siRNA to a final concentration of 25 µM with Opti-MEM (Thermo Fisher Scientific) and Lipofectamine 2000 Transfection Reagent (Thermo Fisher Scientific). All siRNAs were purchased from Dharmacon (Lafayette, CO). After 24h of transfection, the media containing siRNA were removed, and the cells were treated with either 0.1% ETOH or 10nM R5020 for another 72h. The cells were then collected for further analysis, such as flow cytometry and Western blotting analysis.

### 2.10. Analysis of Cytochrome C release by Flow Cytometry

MCF-7PR cells were plated at a density of 3 × 10^5^ cells in 60 mm culture dish. After 48h of starvation in phenol red-free media, the cells were treated with 0.1% ETOH or 10 nM R5020 for 72h, followed by subculture for another 24, 48, or 72 hours. Both floating and adherent cells were then collected at the specified time points into a 1.5 ml microcentrifuge tube and pelleted by centrifugation at 1,000 × g for 5 minutes. The cell pellets were treated with 200 µl of digitonin (50 µg/ml in PBS with 100 mM KCl) on ice for 3 minutes. The cells were washed with 1 ml of PBS and pelleted at 1,000 × g for 5 minutes. The resulting cell pellets were then resuspended with 4% PFA in PBS and incubated for 20 minutes at room temperature. After incubation, the cells were washed twice with PBS. From this point onwards, the centrifuge speed was set at 4,500 × g and spun for 5 minutes to pellet the cells. The cells were incubated in blocking buffer (3% BSA, 0.05% saponin in PBS) for 1h at room temperature, followed by 1:600 anti-cytochrome c antibody (Biolegend, San Diego, California, LOT number #612310) incubation at 4°C overnight in the dark. The cells were washed once with 1 ml of PBS, then resuspended with 300 µl of PBS for flow cytometry (BD FACSymphony A5.2) analysis.

### 2.11. Immunofluorescent Staining

The cells were grown on coverslips and treated for 72h with either 0.1% ethanol or 10 nM R5020. The cells were then rinsed with PBS three times to remove any floating cells. The cells were fixed with 4% PFA for 30 minutes, followed by permeabilization with 0.2% Triton-X for 20 minutes after rinsing three times to remove excess PFA. Triton-X was then rinsed away, and the cells were blocked with 5% BSA in TBST (Tris-Buffered Saline with Tween 20) for 1h. Primary antibodies for TOMM20 (Santa Cruz Biotechnology Inc., Dallas, TX, USA, LOT number sc-17764 PE, 1:100), BNIP3L/NIX (Cell Signaling Technology, Danvers, MA, USA, LOT number #12396, 1:200), Alexa Fluor 647-conjugated cytochrome c antibody (Biolegend, San Diego, California, LOT number #612310, 1:200), BAX (Cell Signaling Technology, Danvers, MA, USA, LOT number #2772, 1:200), and BAK (Santa Cruz Biotechnology Inc., Dallas, TX, USA, LOT number #517390, 1:200) were incubated with the cells overnight. After overnight incubation, the cells were washed three times, with each wash lasting 10 minutes. The cells were then stained with secondary antibodies (1:1000) conjugated with Alexa Fluor 555 (Cell Signaling Technology, Danvers, MA, USA, LOT number #4413) or FITC (Thermo Scientific, Rockford, IL, USA, LOT number #31569) for 1h at room temperature (except for the Alexa Fluor 647-conjugated cytochrome c). The cells were then counterstained with DAPI after washing three times with PBS and mounted with antifade mountant on microscope slides (Thermo Scientific, Rockford, IL, USA, LOT number #P36930). The images were visualized and acquired using a fluorescent microscope (Zeiss). Post-processing of images was done using Zen software.

### 2.12. JC-1 staining

The cells were treated for 72+48 hr or 72+72 hr for mitochondrial membrane potential measurement using JC-1 dye. The staining solution was prepared in the culture media, with 2 µM as the final concentration of JC-1 dye (MedChemExpress, USA). Both adherent and floating cells were stained with JC-1 dye for 15 minutes before being trypsinized and pelleted by centrifugation. The cells were then resuspended in 1X PBS and analysed by flow cytometry (BD FACSymphony A5.2).

### 2.13 MitoTracker Staining

The cells were plated on a glass-bottom 35 mm culture dish and treated for 72h+72h with either 0.1% ETOH or 10 nM R5020. Before staining, the cells were rinsed once with DPBS, followed by the addition of media containing MitoTracker™ Red CMXRos Dye (Invitrogen, Carlsbad, CA, USA, LOT number #M7512), using 20 nM as the final concentration. After a 20-minute incubation, the media was removed and replaced with fresh media before proceeding with observation under a fluorescence microscope.

### 2.14. Protein Lysate Collection and Western Blotting Analysis

Cells were lysed in cold lysis buffer [50 mM HEPES-KOH (pH 7.5), 100 mM NaF, 150 mM NaCl, and 1% Triton X-100], and the supernatant was collected after centrifugation. Proteins in the total cell lysates were quantified, resolved by SDS-PAGE electrophoresis, and transferred onto a PVDF membrane. After blocking with 2.5% BSA in Tris-buffered saline with Tween 20 (TBST), the membranes were incubated with primary antibodies overnight at 4 °C. The primary antibodies used in the experiments are listed in the table below. The secondary antibodies, anti-mouse (1:1000) and anti-rabbit (1:1000) (Cell Signaling Technology, Danvers, MA, USA), conjugated with horseradish peroxidase (HRP), were used according to the source of the primary antibodies. Proteins on the membranes were detected using Immobilon Western Chemiluminescent HRP substrate (Merck Millipore, Billerica, MA, USA) and the ChemiDoc MP Imaging System (Bio-Rad Laboratories, Inc.).

| Antibody | Dilution | Source | Catalogue No. |
| --- | --- | --- | --- |
| H190 Total PR | 1:1000 | Santa Cruz Biotechnology Inc., Dallas, TX, USA | sc-7208 |
| ER-α | 1:1000 | Santa Cruz Biotechnology Inc., Dallas, TX, USA | sc-8002 |
| Caspase 9 | 1:1000 | Santa Cruz Biotechnology Inc., Dallas, TX, USA | sc-56076 |
| Cleaved Caspase 9 | 1:1000 | Cell Signaling Technology, Danvers, MA, USA | 9505 |
| Caspase 7 | 1:1000 | Cell Signaling Technology, Danvers, MA, USA | 9492 |
| Caspase 8 | 1:1000 | Cell Signaling Technology, Danvers, MA, USA | 4790 |
| BID | 1:1000 | Cell Signaling Technology, Danvers, MA, USA | 2022 |
| BAX | 1:1000 | Cell Signaling Technology, Danvers, MA, USA | 2772 |
| PARP | 1:1000 | Cell Signaling Technology, Danvers, MA, USA | 9542 |
| BAK | 1:1000 | Cell Signaling Technology, Danvers, MA, USA | 3814 |
| BCL-2 | 1:1000 | Santa Cruz Biotechnology Inc., Dallas, TX, USA | sc-7382 |
| BCL-XL | 1:1000 | Santa Cruz Biotechnology Inc., Dallas, TX, USA | Sc-8392 |
| HIF1A | 1:1000 | Cell Signaling Technology, Danvers, MA, USA | 36169 |
| BNIP3 | 1:1000 | Cell Signaling Technology, Danvers, MA, USA | 44060 |
| BNIP3L/NIX | 1:1000 | Cell Signaling Technology, Danvers, MA, USA | 12396 |
| LC3B | 1:1000 | Cell Signaling Technology, Danvers, MA, USA | 83506 |
| HTRA2/OMI | 1:1000 | Cell Signaling Technology, Danvers, MA, USA | 9745 |
| Endonuclease G | 1:1000 | Cell Signaling Technology, Danvers, MA, USA | 4969 |
| TNFAIP3 | 1:1000 | Cell Signaling Technology, Danvers, MA, USA | sc-166692 |
| BIRC3 | 1:1000 | Cell Signaling Technology, Danvers, MA, USA | 3130 |
| DNA-PKcs | 1:1000 | Cell Signaling Technology, Danvers, MA, USA | 4602 |
| CDK2 | 1:1000 | Santa Cruz Biotechnology Inc., Dallas, TX, USA | 6248 |
| CDK4 | 1:1000 | Santa Cruz Biotechnology Inc., Dallas, TX, USA | sc-23896 |
| Total H1.3 | 1:1000 | Bio-Rad Laboratories, Hercules, California | VPA00783 |
| Phos-H1.3 | 1:1000 | Invitrogen, Carlsbad, CA, USA |  |
| β-actin | 1:1000 | Santa Cruz Biotechnology Inc., Dallas, TX, USA | sc-47778 |
| GAPDH | 1:10000 | Santa Cruz Biotechnology Inc., Dallas, TX, USA | sc-47724 |
| Anti-mouse secondary | 1:1000 | Cell Signaling Technology, Danvers, MA, USA | 7076 |
| Anti-rabbit secondary | 1:1000 | Cell Signaling Technology, Danvers, MA, USA | 7074 |

