## Supplementary Figure 1 - 4 for "Omics analysis reveals striking effects of progesterone receptor on mitochondria and mitochondria-mediated apoptosis independent of caspases in Breast Cancer cells"

### Supplementary Fig. 1

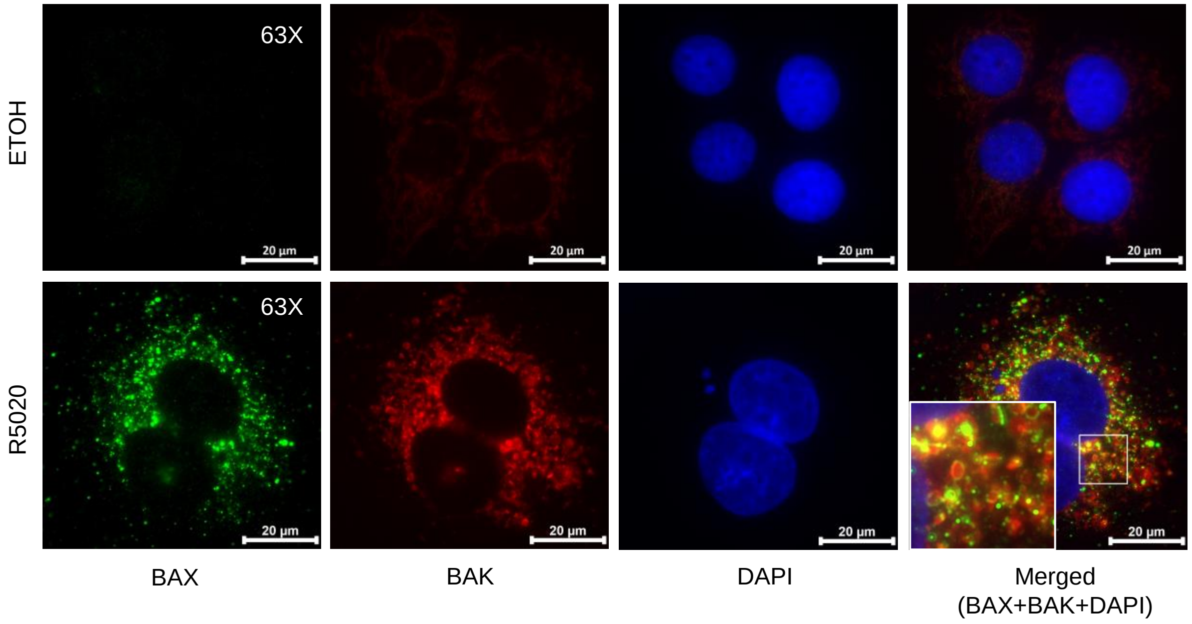


**Supplementary Fig. 1 Agonist-activated PR induced BAX and BAK punctate formation.** Co-immunostaining of BAX, BAK, and DAPI was performed on MCF-7PR cells treated with control or R5020 for 72h. In the control cells, the composite image shows only DAPI (blue), as no BAX or BAK was observed. Cells treated with R5020 displayed punctate BAX (green) and BAK (red), with some overlap in the composite image (yellow). This corresponds to the understanding that BAX and BAK can form pores independently or through hetero-oligomerization.

### Supplementary Fig. 2

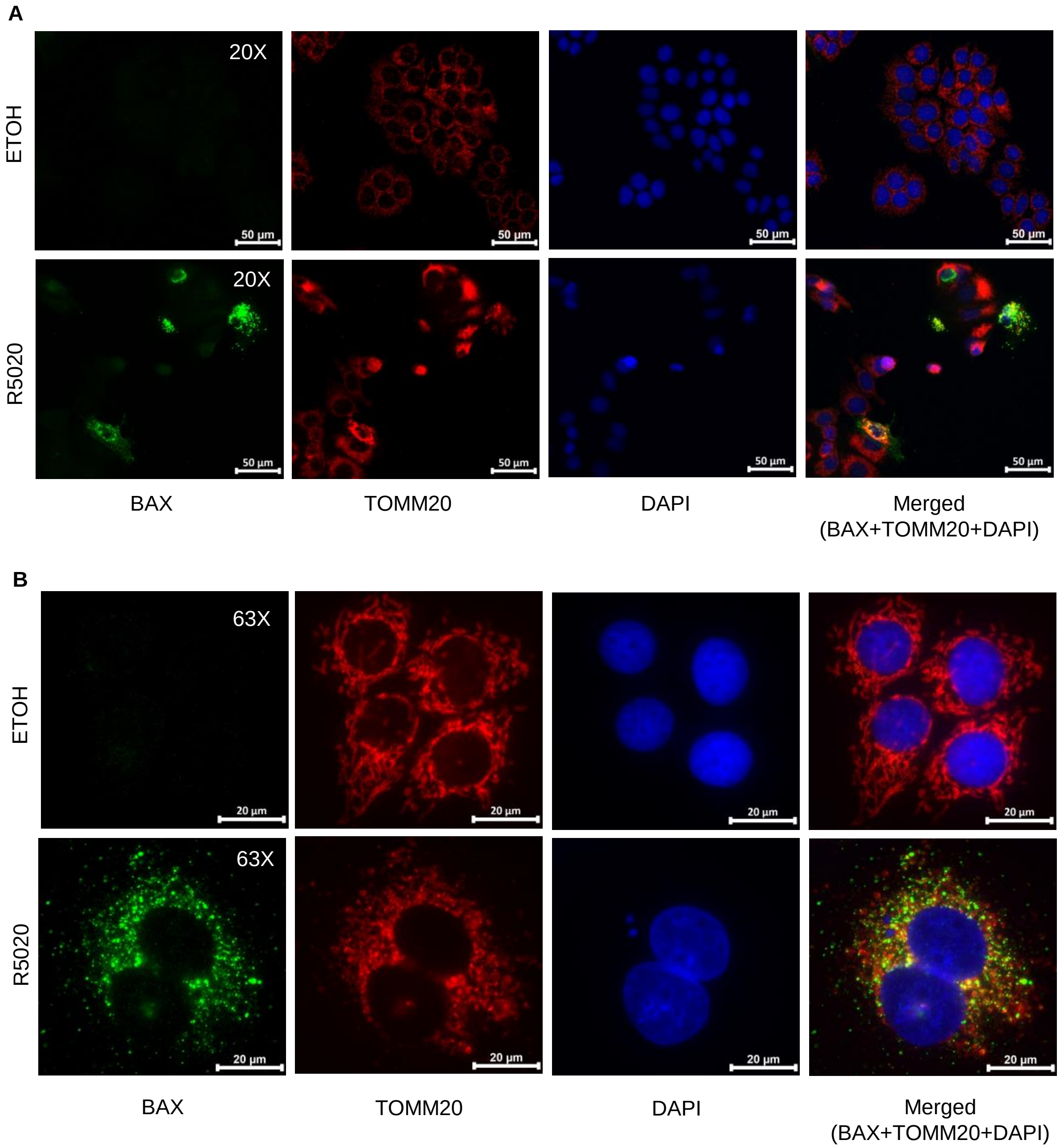


**Supplementary Fig. 2 Agonist-activated PR induced BAX translocation to the mitochondrial outer membrane.** Co-immunostaining of BAX, TOMM20, and DAPI was performed on MCF-7PR cells treated with control or R5020 for 72h + 72h. **(A)** A 20X objective lens was used. BAX was undetectable in control cells, while it was observed in a few cells treated with R5020. The overlap of BAX (green) and TOMM20 (red) is visible in the composite image, as activated BAX is known to translocate to the mitochondrial outer membrane. **(B)** At higher magnification (63X), BAX appeared as clustered punctate structures, either fully, partially, or adjacent to the mitochondrial staining in R5020-treated cells. The mitochondrial marker (TOMM20) appeared as an interconnected, network-like structure, evenly distributed around the nucleus (DAPI) in control cells. However, R5020 treatment altered the structure into spherical aggregates.

### Supplementary Fig. 3

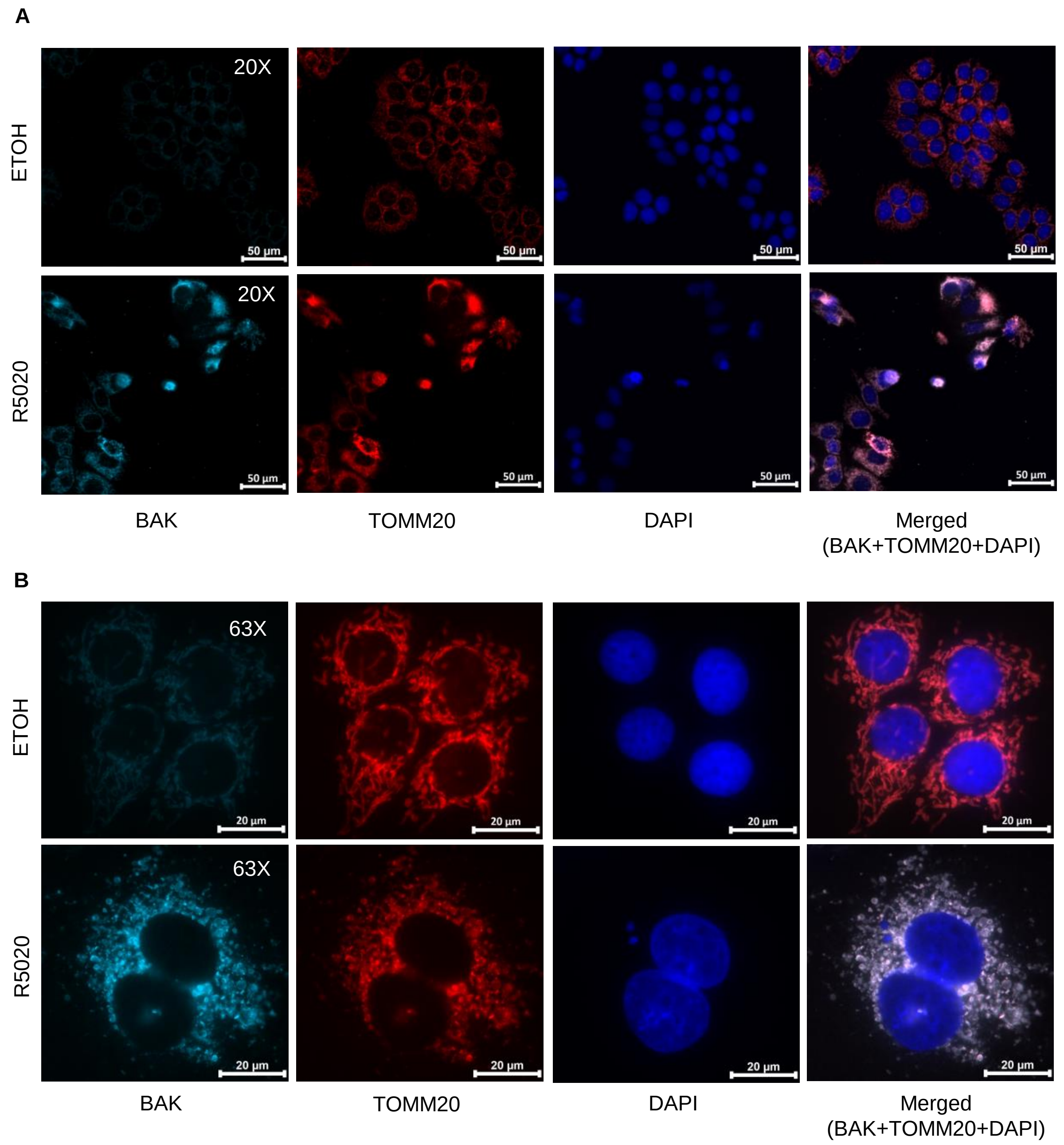


**Supplementary Fig. 3 Agonist-activated PR increased BAK immunostaining in the mitochondria.** Co-immunostaining of BAK, TOMM20, and DAPI was performed on MCF-7PR cells treated with control or R5020 for 72h. **(A)** A 20X objective lens was used. BAK was undetectable in control cells, so only TOMM20 (red) and DAPI were shown in the composite image. In contrast, BAK (light blue) was observed in all cells treated with R5020 and completely overlapped with the mitochondria, resulting in a light pink color after merging with the mitochondria marker (TOMM20). **(B)** At 63X magnification, the images clearly showed that BAK was largely colocalized with the mitochondrial staining, with many punctate structures observed in R5020-treated cells undergoing mitochondrial fragmentation and morphological changes.

### Supplementary Fig. 4

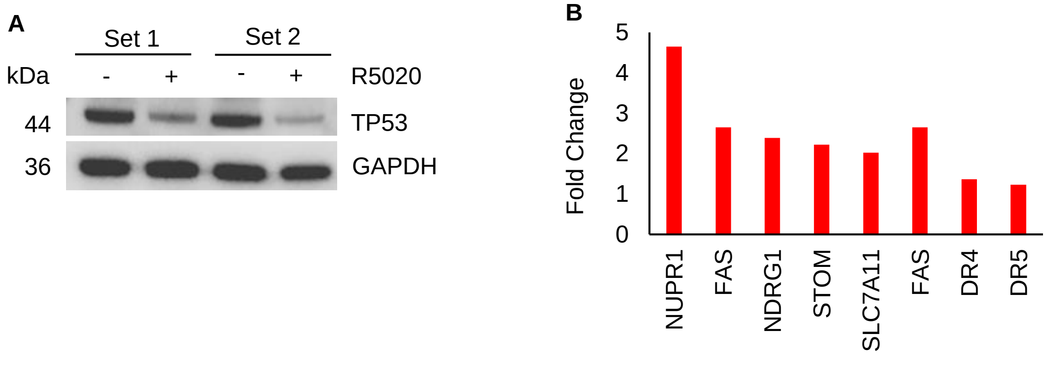


**Supplementary Fig. 4 The p53 pathway is also involved in agonist-activated PR-induced apoptosis. (A)** Downregulation of TP53 by R5020 treatment was detected, despite not being observed in the MS data. The downregulation of p53 protein may result from increased p53 transcriptional activity due to DNA binding, which enhances its ubiquitination, or from the MDM2 negative feedback mechanism, even though the p53 pathway was upregulated by R5020 treatment. GAPDH was used as a loading control. **(B)** The top 5 upregulated proteins (NUPR1, FAS, NDRG1, STOM, SLC7A11) and death receptors (DR4 and DR5) involved in the p53 pathway hallmark are shown in the bar plot. The red bars indicate upregulation of the proteins in the MS results.
